# A 19-isolate reference-quality global pangenome for the fungal wheat pathogen *Zymoseptoria tritici*

**DOI:** 10.1101/803098

**Authors:** Thomas Badet, Ursula Oggenfuss, Leen Abraham, Bruce A. McDonald, Daniel Croll

## Abstract

**Background:** The gene content of a species largely governs its ecological interactions and adaptive potential. A species is therefore defined by both core genes shared between all individuals and accessory genes segregating presence-absence variation. There is growing evidence that eukaryotes, similar to bacteria, show intra-specific variability in gene content. However, it remains largely unknown how functionally relevant such a pangenome structure is for eukaryotes and what mechanisms underlie the emergence of highly polymorphic genome structures.

**Results:** Here, we establish a reference-quality pangenome of a fungal pathogen of wheat based on 19 complete genomes from isolates sampled across six continents. *Zymoseptoria tritici* causes substantial worldwide losses to wheat production due to rapidly evolved tolerance to fungicides and evasion of host resistance. We performed transcriptome-assisted annotations of each genome to construct a global pangenome. Major chromosomal rearrangements are segregating within the species and underlie extensive gene presence-absence variation. Conserved orthogroups account for only ∼60% of the species pangenome. Investigating gene functions, we find that the accessory genome is enriched for pathogenesis-related functions and encodes genes involved in metabolite production, host tissue degradation and manipulation of the immune system. *De novo* transposon annotation of the 19 complete genomes shows that the highly diverse chromosomal structure is tightly associated with transposable elements content. Furthermore, transposable element expansions likely underlie recent genome expansions within the species.

**Conclusions:** Taken together, our work establishes a highly complex eukaryotic pangenome providing an unprecedented toolbox to study how pangenome structure impacts crop-pathogen interactions.

## Background

Microbial species harbor substantial functional diversity at the level of gene presence-absence variation (1). Genes not fixed within a species (*i.e.* accessory genes) can account for a large fraction of the full gene repertoire (*i.e.* the pangenome). In bacteria, the proportion of core genes in the pangenome can range from 5-98% and challenge taxonomic classifications (2, 3). The wide spectrum of pangenome sizes across species can be associated with the species distribution and lifestyle (4). Species showing a wide geographical distribution and large population sizes characterized by frequent genetic exchange tend to have expansive, open pangenomes (5). In microbial pathogens, accessory genes play a major role in virulence and environmental adaptation (6–8). The notion of a pangenome led to the discovery that major elements of intra-specific variation are often ignored in studies relying on a single reference genome. Large pangenomes also can challenge association studies aiming to identify the genetic basis of phenotypic traits because mapping is often performed against a single reference genome, making potentially relevant genetic variation inaccessible (9, 10). Despite their importance for unravelling the genetic basis of adaptive evolution, only a very limited number of eukaryotic species have well established pangenomes.

Copy number variation including gene deletion generates intraspecific gene content variation in nearly all species (11). This variation can create extreme variance in fitness and promote adaptive evolution (12–15). In plant pathogens, the ability to infect a host often relies on the secretion of effector proteins that interfere with the host cell machinery (16–18). Host plants evolved cognate resistance proteins that are able to recognize effector proteins and trigger immunity (19). Gains and losses of effector genes can therefore have a major impact on the outcome of host-pathogen interactions and challenge food security. Recent studies on fungal pathogens highlighted that genes showing presence-absence variation are enriched for predicted effectors (14,20,21). Effectors and transposable elements (TEs) are often tightly associated with fast-evolving compartments of the genome (22, 23), also known as the “two-speed” genome architecture (24). However, how TEs impact the birth and death of effectors in fast-evolving compartments remains largely unclear (6, 25). The construction of pathogen pangenomes enabled crucial insights into functional diversity and the evolutionary trajectories of host adaptation. Recent pangenome analyses of four fungal species including opportunistic pathogens revealed that between ∼9-19% of the pangenome is accessory. Accessory gene localization was preferentially in subtelomeric regions, suggesting both a mechanistic link to repeat-rich regions and relaxation of selective constraints (26). The wheat pathogen *Zymoseptoria tritici* was found to have one of the largest eukaryotic pangenomes with an estimate of at least 42% of all genes being accessory (27). However, eukaryotic pangenomes remain shallow and are often based on not fully resolved chromosomal sequences.

Fungal plant pathogens such as *Z. tritici* show extreme cases of genome plasticity. The reference genome of *Z. tritici* has 21 chromosomes, of which eight are accessory and segregate presence-absence variation in populations (28). The pathogen rapidly evolved virulence on resistant wheat cultivars and has overcome all current fungicides (29–31). Host adaptation was driven among other factors by the rapid deletion of an effector gene and structural rearrangements (32–34). Pathogen populations are highly diverse with high rates of recombination (35–37). Meiosis can trigger large chromosomal rearrangements and lead to aneuploid chromosomes in the species (38, 39). A pangenome constructed for five *Z. tritici* isolates revealed that chromosome length variation segregating within populations was mainly due to the presence-absence variation of large TE clusters (27, 40). Furthermore, accessory genes tended to form clusters dispersed along chromosomes. Accessory genes also tended to be in closer proximity to TEs than core genes and were therefore more likely to be affected by epigenetic silencing (27). However, the constructed pangenome was very likely incomplete given the fact that four of the genomes originated from isolates collected in the same year from two nearby fields. Furthermore, accessory genes were enriched for pathogenesis-related functions but the pangenome size did not reach saturation. Given the global impact of the pathogen and the importance of accessory genes for adaptive evolution, a comprehensive pangenome capturing worldwide genetic diversity is essential.

In this study, we construct the pangenome of *Z. tritici* by including 19 isolates sampled from six different continents and covering the global distribution of the pathogen. We test to what extent the species segregates chromosomal rearrangements and how this impacts gene presence-absence variation at loci relevant for pathogenicity. We also analyze whether TE content is polymorphic within the species and may contribute to genome size evolution.

## Results

### Major chromosomal rearrangements segregating within the species

We constructed a global pangenome of *Z. tritici* based on 19 isolates sampled from six continents and 13 different countries (Figure 1A-B). The isolates included the previously described reference isolate IPO323 sampled in the Netherlands and four isolates that were isolated from two nearby fields in Switzerland (27,28,40). The geographic regions of origin of the 19 isolates recapitulate a significant environmental gradient in mean annual temperature and humidity and span the distribution range of the species. The sampling period ranges from 1984 (IPO323) to 2010 (CRI10). Fungicide applications against *Z. tritici* became widespread in the 90s and early 2000s, hence the sampling covers both pre- and post-fungicide treatment regimes. We sequenced long-read PacBio SMRTbell libraries to a depth of 40-110X and ∼20 kb read coverage in order to generate chromosome-level assemblies. Assembly sizes ranged from 37.13 Mb (IR01_48b) to 41.76 Mb (Aus01) (Figure 1C).

**Figure 1:**
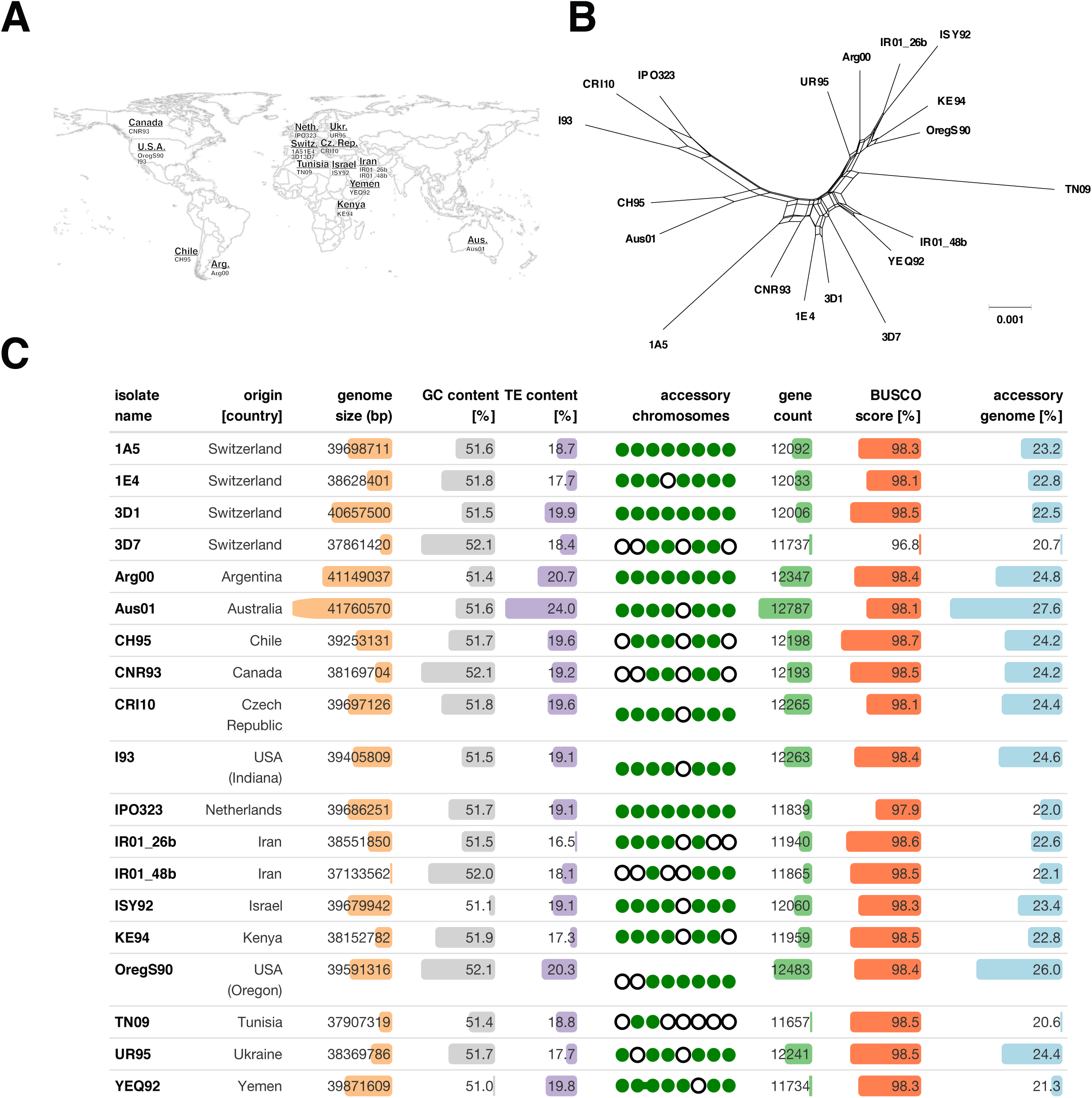
Assembly of 19 complete genomes from a worldwide collection. **A**. World map indicating the isolate names and country of origin. **B**. Phylogenomic tree based on 50 single-copy orthologs showing reticulation using SplitsTree. **C**. Summary of genome assembly characteristics for all isolates. The bars represent the range of minimum (shortest bar) to maximum values (longest bar) for each reported statistic. Chromosome 14-21 are accessory chromosomes. The presence or absence of accessory chromosomes in each genome is shown by green dots and empty circles for present and missing chromosomes, respectively. The linked dots for isolate YEQ92 indicate the chromosomal fusion event (see also Figure 2).

We recovered all eight known accessory chromosomes of the species but no additional chromosome. The accessory chromosome 18 is most often missing. Together, the 8 accessory chromosomes display an average size variation of ∼37% across all isolates and a maximum of 60% for chromosome 14 (Figure 2A). For core chromosomes, the average size variation accounts for 16% of chromosome length going up to 23% for chromosome 7. We identified a major deletion spanning 406 kb and encompassing 107 genes on the right arm of core chromosome 7 of the Yemeni isolate (YEQ92; Figure 2B lower panel). The same isolate had chromosome 15 fused to the right arm of chromosome 16. The fusion event is supported by aligned PacBio reads spanning the region between the two chromosomal segments (Additional file 1: Figure S1). The resulting chromosome is 1.20 Mb long and 49.5 kb shorter than the sum of the homologous chromosomes 15 and 16 of the IPO323 reference genome. Approximately 90% of the genes on the IPO323 chromosome 15 and 16 belong to accessory orthogroups, as they lack an ortholog in at least one of the other isolates. We find that the chromosomal fusion deleted about 150 kb affecting 1 and 12 genes on chromosomes 15 and 16, respectively (Figure 2B upper panel). We further assessed genome completeness using BUSCO analyses. All genomes exceed the completeness of the fully finished IPO323 reference genome (97.9%) with the exception of isolate 3D7 (96.8%; Figure 1C).

**Figure 2:**
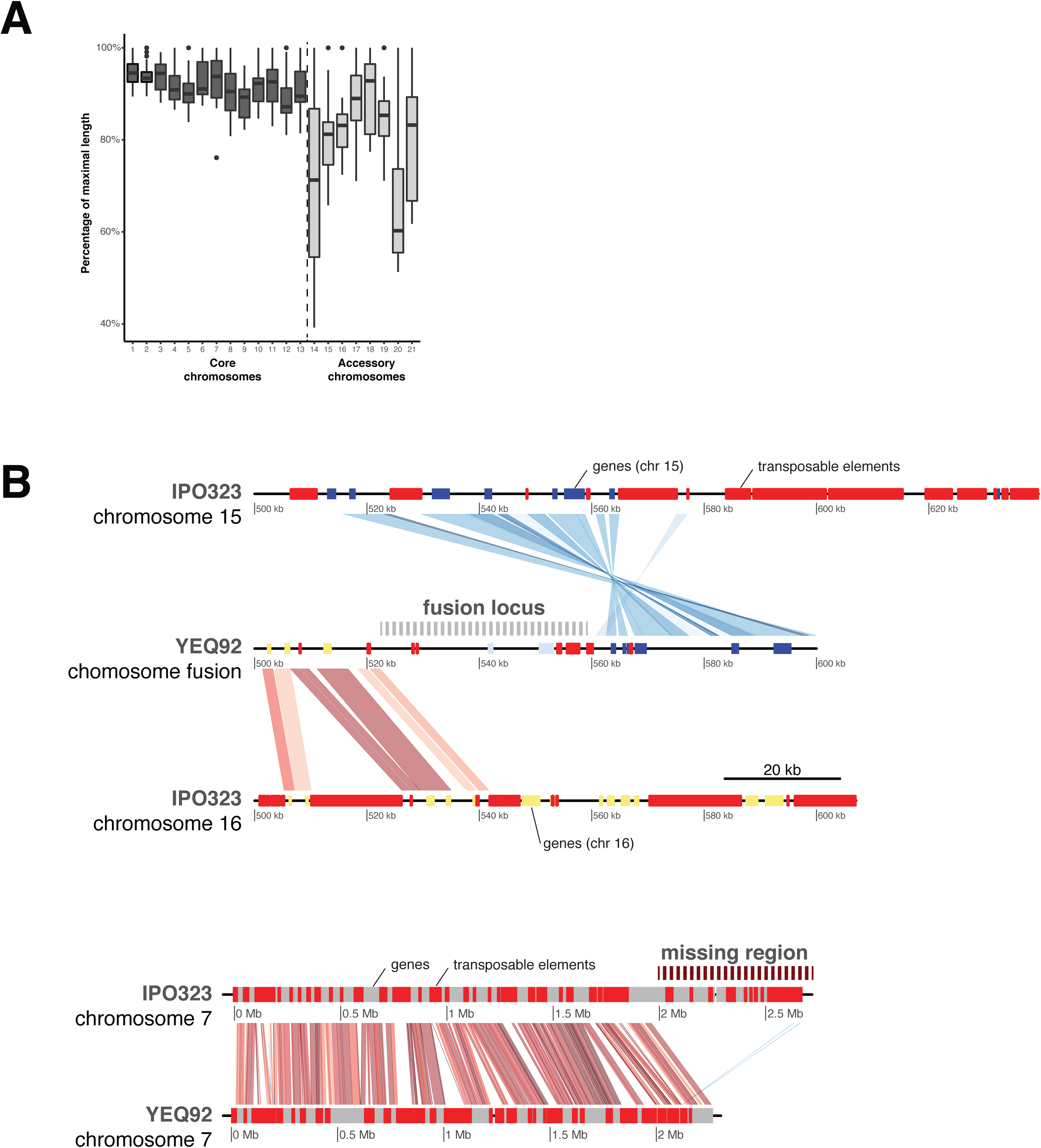
Large segregating chromosomal rearrangements within the species. **A.** Chromosome length variation expressed as the percentage of the maximum observed length for each chromosome. **B.** Two large chromosomal rearrangements identified in the isolate YEQ92 isolated from Yemen. The upper part shows the local chromosomal synteny at the fusion locus between accessory chromosomes 15 and 16 identified in YEQ92 compared to the reference genome IPO323. Transposons are shown in red, genes from chromosome 15 in purple, genes from chromosome 16 in green and genes specific to the fusion in grey boxes, respectively. Synteny shared between chromosomes is shown in red for colinear blocks or blue for inversions. The lower part shows the whole chromosome synteny of chromosome 7 contrasting YEQ92 to the reference genome IPO323. YEQ92 misses a subtelomeric region. Transposons are shown in red and genes in grey.

### Substantial gene content variation across the pangenome

We generated RNAseq data to identify high-confidence gene models in all 14 newly assembled genomes based on a splice-site informed gene prediction pipeline. The total gene count varied between 11’657 and 12’787 gene models (Figure 1C). We assigned all genes to orthogroups using protein homology and constructed a pangenome of all 19 complete genomes. The pangenome consists of a total of 229’699 genes assigned to 15’474 orthogroups. The number of genes assigned per orthogroup varies among isolates (Figure 2B). Approximately 99.8% of all orthogroups (15’451) are single gene orthogroups and ∼60% of all orthogroups are shared among all 19 isolates (9’193 core orthogroups). Around 96% of the core orthogroups (8’829 out of 9’193) have conserved gene copy numbers among isolates. Furthermore, we find that 30% of all orthogroups are shared between some but not all genomes (4’690 accessory orthogroups) and 10% of the orthogroups are composed of genes found in a single genome only (1’592 singletons; Figure 3A-B; Additional file 2: Table S1).

**Figure 3:**
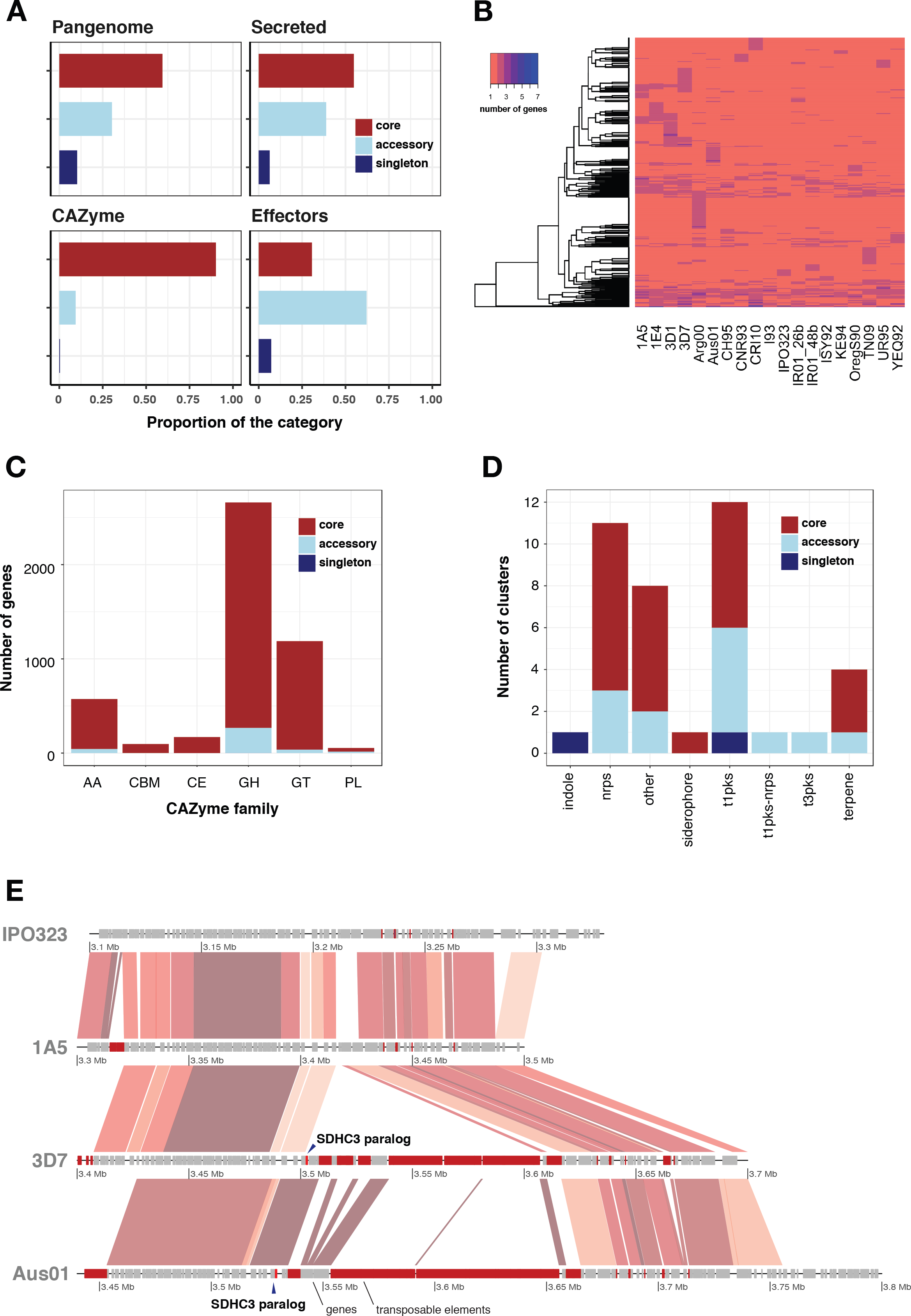
Construction and analysis of the *Zymoseptoria tritici* pangenome. **A**. Proportions of core orthogroups (present in all isolates), accessory orthogroups (present ≥ 2 isolates but not all) and singletons (present in one isolate only) across the pangenome (upper-left). The proportions of core, accessory and singleton categories are shown for orthogroups coding for secreted proteins (upper-right), carbohydrate-active enzymes (CAZymes; lower-left) and effectors (lower-right). **B**. Gene copy number variation in core orthogroups across the 19 genomes. **C**. Pangenome gene count across six CAZyme families. Families are divided into glycoside hydrolase (GH), glycosyl transferase (GT), auxiliary activity (AA), carbohydrate esterase (CE), carbohydrate-binding modules (CBM) and polysaccharide lyase activity (PL) categories. **D**. Pangenome categories of secondary metabolite gene clusters. **E**. Synteny plot of succinate dehydrogenase (SDH) paralogs mediating fungicide resistance. The SDHC3 locus on chromosome 3 is shown for isolates 3D7 and Aus01 both carrying the paralog. IPO323 and 1A5 lack SDHC3. The position of the SDHC3 paralog is shown using dark arrows. Genes are colored in grey and transposable elements in red.

To infect wheat, *Z. tritici* relies on specific gene functions (41, 42). Effectors play a major role in establishing infection and exploiting host resources. Hence, we analysed how gene functions were structured across the pangenome components. Core orthogroups showing variation in gene-copy number among isolates include five encoding predicted effectors. Both accessory proteins and overall effector proteins are less conserved than core proteins at the amino acid level (Additional file 1: Figure S2). A total of 3.5% (691) of all orthogroups encode at least one predicted effector. Among orthogroups encoding at least one predicted effector, 31% were conserved among all isolates (219), 63% were accessory (436) and 5% were found in only one isolate (36 singletons). Notably, 99% of the predicted effector genes are located on core chromosomes. In addition to effectors, enzymes enabling access to nutrients are important pathogenicity components. We identified a total of 4’742 annotated carbohydrate-degrading enzymes (CAZymes) clustered into 263 orthogroups. Notably, 92% of the orthogroups encoding CAZymes were conserved among all isolates (Figure 3A). CAZymes grouped into 123 subfamilies. Glycoside hydrolases (GH) are the largest family and account for 57% of all annotated CAZymes (151 orthogroups for 2’717 genes). Glycosyl transferases (GT) are the second most abundant family with 1’188 genes and 66 orthogroups (25% of all CAZymes) (Figure 3C). We also identified 33 orthogroups encoding for auxiliary activities (AA), 9 for carbohydrate esterase activity (CE), 6 for carbohydrate-binding modules (CBM) and 3 for polysaccharide lyase activity (PL). The PL family includes 29% accessory genes. Across CAZyme families, 0-10% of the genes are accessory (Figure 3C). We found a singleton GH43 subfamily gene in the genome of the Australian isolate (Aus01).

The production of secondary metabolites contributes significantly to virulence and competitive abilities of fungal pathogens. We identified between 29 and 33 secondary metabolite gene clusters per genome depending on the isolate. A total of 70% of all genes predicted as components of a biosynthetic gene cluster are conserved between all isolates and 30% are accessory (Figure 3D, Additional file 1: Figure S3). Of the 147 orthogroups annotated as encoding biosynthetic or biosynthetic-additional proteins in the pangenome, 87, 92, 111 and 112 have a homolog with >50% identity in the four closely related sister species *Z. passerinii*, *Z. ardabiliae*, *Z. pseudotritici* and *Z. brevis*, respectively (Additional file 1: Figure S4). We identified 39 syntenic gene clusters in the pangenome classified into 12 type 1-polyketide synthase (PKS), 11 non-ribosomal peptide synthetase (NRPS), four terpene, one type 3-PKS, one siderophore, one indole and eight unclassified clusters. Sixteen (40%) of the identified syntenic clusters show presence-absence variation. In the CH95 isolate, a gene cluster on chromosome 7 was annotated as unclassified but annotated as a NRPS in 17 other isolates and absent from the IPO323 reference genome. The sole indole and type 1-PKS clusters located on chromosomes 5 and 10, respectively, were only found in isolate TN09. Two type 1-PKS and one NRPS cluster were missing in the isolates YEQ95, Aus01 and IPO323, respectively. Among the 39 identified syntenic gene clusters, 23 included a predicted effector and nine included a gene annotated as a cell-wall degrading enzyme.

The emergence of fungicide tolerance in *Z. tritici* is a major threat to wheat production. Succinate dehydrogenase (SDH) inhibitors are commonly used as control agents (31, 43). We identified five SDH orthologs, of which three were conserved among all genomes (SDHB, SDHC and SDHD subunits). We find two distinct SDHC paralogs SDHC2 and SDHC3 in eleven and two isolates, respectively. The SDHC3 paralog conferring standing resistance to SDH inhibitors is located adjacent to a large cluster of TEs, suggesting that chromosomal rearrangements were underlying the paralog emergence (Figure 3E). Genes encoding major facilitator superfamily (MFS) transporters, which can confer multidrug resistance in *Z. tritici* (44), grouped into 336 orthogroups for a total of 5’787 genes (Additional file 2: Table S2). We find that 39 (11%) of these orthogroups are part of a predicted secondary metabolite gene cluster and one is an annotated CAZyme from the GH78 family. Overall, the results reveal that gene families essential for pathogenicity and fungicide resistance show unexpectedly high levels of presence-absence variation in the *Z. tritici* pangenome.

### Strong expression variation across major gene functions

Differential gene expression is a major driver of intraspecific phenotypic differences. We performed mRNA-sequencing of all 19 isolates grown on minimal media. Minimal media induces filamentous growth of *Z. tritici*, mimicking the morphology and nutrient starvation that occurs early during plant infection. We investigated isolate-specific gene expression by self-mapping RNAseq reads to each isolate’s genome assembly. Overall, 91.3% of the genes show expression on minimal media and 68% have expression of more than 10 counts per million (CPM) (Figure 4A). Core genes have higher expression than accessory genes (Additional file 1: Figure S5). Among the genes showing no expression on minimal media, 501 are predicted effector genes (8% of predicted effectors), 93 are predicted CAZymes (2% of CAZymes) and 838 are members of a predicted gene cluster (10% of all gene cluster genes). CAZymes are overall highly expressed on minimal media (∼77% with CPM >10) when compared to effectors (∼45% with CPM >10) and gene cluster genes (∼60% with CPM >10) (Figure 4A). About 53% of core single copy orthogroups with non-zero expression have a coefficient of variation >50% (Figure 4B). Similarly, ∼68% of CAZymes and ∼60% of genes that are part of a secondary metabolite cluster have expression coefficient of variation > 50%. In contrast, about 90% of orthogroups encoding predicted effectors have a coefficient of variation >50%, together with ∼81% of accessory orthogroups.

**Figure 4:**
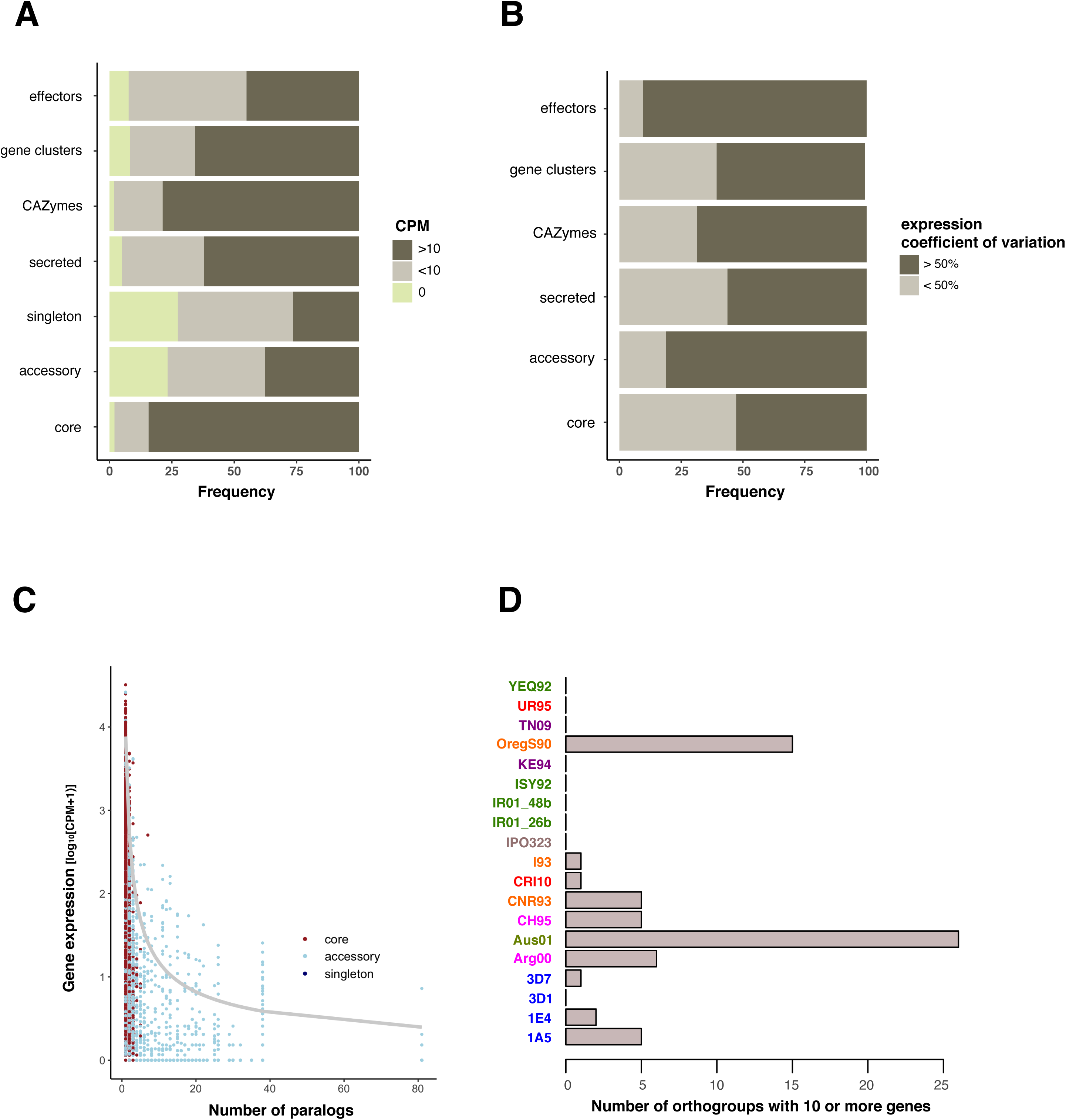
Expression polymorphism across the pangenome. **A.** Proportion of genes showing expression >10 counts per million (CPM) across genes categories. The frequencies are shown for orthogroups encoding putative effectors, secondary metabolite cluster genes (gene cluster), carbohydrate-active enzymes (CAZymes), secreted proteins. The frequencies are also shown for singleton, accessory and core orthogroup categories in the pangenome. **B**. Proportion of orthogroups for which the expression coefficient of variation is >50% [*cov* = *sd* (CPM) / *mean* (CPM)] among different gene and pangenome categories as in (A). **C**. Correlation of gene expression and the number of paralogs detected for the same gene per genome. The grey line shows the logarithmic regression based on the linear model *log_10_* (CPM+1) *∼ log_10_* (number of paralogs). **D**. Number of orthogroups with ≥10 paralogs per genome. Isolates are colored by continent of origin.

To identify broad patterns in the pangenome expression landscape, we performed a clustering analysis of all core single gene orthogroups. We find that expression clustering does not reflect the geographical origin or genetic distance with the exception of the four Swiss isolates (1A5, 1E4, 3D1 and 3D7; Additional file 1: Figure S6). We also analysed the impact of copy-number variation on average expression and find that single-copy orthologs are on average more highly expressed. In addition, we show that gene expression rapidly decreases if an orthogroup includes 2-8 paralogs (Figure 4C).

### A highly variable transposable element content within the species

TEs are drivers of pathogen evolution by generating adaptive genetic variation. To identify genes with a potential role in the mobilisation of TEs, we analysed large homology groups. Among the orthogroups with 10 or more paralogs, ∼88% of the genes encode proteins without homology in databases, ∼7% of the genes encode nucleic acid binding functions (GO:0003676), ∼2% of the genes encode a retrotransposon nucleocapsid (GO:0000943) and ∼1.5% of the genes encode a DNA integration domain (GO:0015074). Orthogroups with 10 or more paralogs are all accessory. For isolates sharing the same large orthogroups, we identified variability in the gene copy number within those orthogroups. Indeed, the isolates Aus01 and OregS90 have 26 and 16 orthogroups, respectively, with more than 10 assigned genes. The isolates I93 and Arg00 count between one and six orthogroups and nine other isolates have no orthogroups larger than ten genes (Figure 4D). Altogether, these results suggest that large orthogroups (>10 genes) essentially regroup genes that are encoded by TEs. Our data also indicates regional TE-driven genome expansions given the enlarged genome sizes in Australian and North American isolates.

To elucidate the role of transposition on generating genomic variation, we screened the 19 genomes for TE content. For this, we jointly analysed all complete genomes to exhaustively identify repetitive DNA sequences. We identified a total of 304 high-quality TE family consensus sequences grouped into 22 TE superfamilies. The GC-content of the consensus sequences is highly variable, ranging from 23-77% (Additional file 1: Figure S7). On average, TE superfamilies have a GC-content lower than 50%, except for unclassified SINE families (RSX; GC% ∼50.6). The genomic TE content ranges from 16.48% (IR01_26b) to 23.96% (Aus01) and is positively correlated with genome size (cor = 0.78, *p* < 0.001; Figure 5A). Genome size correlates with genome-wide TE proportions on both core and accessory chromosomes but is negatively correlated with the proportion of coding sequences (Additional file 1: Figure S8, S9). The average length of individual TEs ranges from 102 to 51’298 bp with the Helitron superfamily having the higher average length (Additional file 1: Figure S10-S11). The largest element is an unclassified LTR (RLX_LARD_Thrym) on chromosome 7, the size of which ranges from 6’282 bp in CNR93 to 59’390 bp in ISY92. This particular LTR is present at the locus only in 18 isolates including ISY92, which has a fragmented secondary copy on chromosome 3. The RLX_LARD_Thrym insertion on chromosome 7 overlaps with the ribosomal DNA locus and showed far above average mapped PacBio read coverage (∼250X).

**Figure 5:**
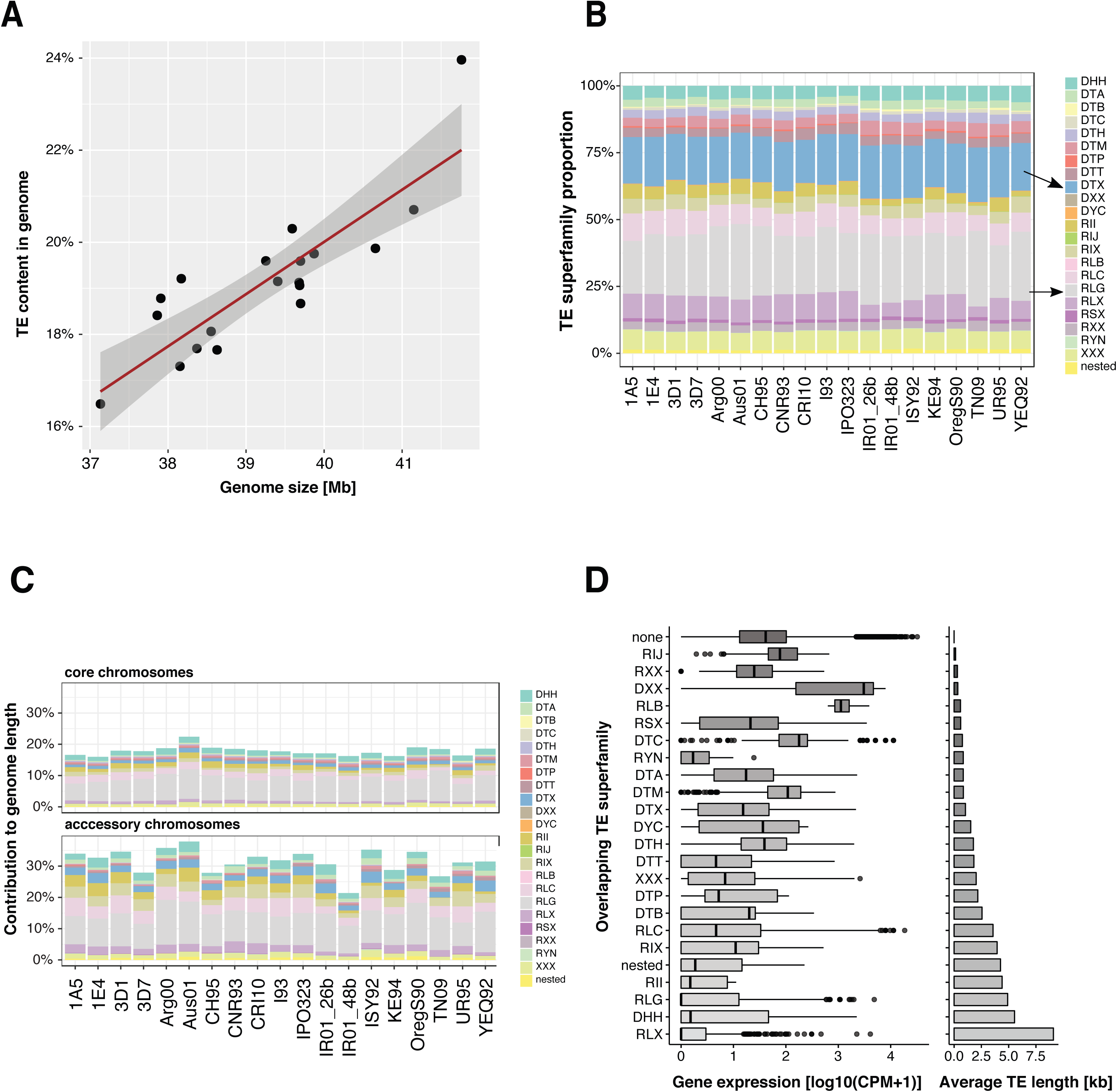
Transposable elements (TEs) and genome size variation. **A**. Contribution of TEs (%) to total genome size across the 19 isolates. **B**. Relative frequency of the 23 TE superfamilies across all genomes with 100% referring to the total TE content of the respective genome. **C**. Contribution of TE superfamilies to core and accessory genome size across the 19 isolates. **D**. Expression of genes affected by TE insertions (grouped by TE superfamilies; left panel) and the mean TE length in the genome (grouped by TE superfamilies; right panel).

The genome-wide content of TEs shows substantial variation among the 19 isolates, however the relative abundance of different TE superfamilies is relatively conserved with LTR *Gypsy*, unclassified TIR and LTR *Copia* elements being the most frequent (Figure 5B). Accessory chromosomes contain consistently higher proportions of TEs compared to core chromosomes (26-41% versus 17-24%; Figure 5C). Aus01 and OregS90 isolates showed the highest TE content. Interestingly, the Aus01 genome shows LINE *I*, LTR *Gypsy* and LTR *Copia* family-specific expansion compared to other genomes. In contrast, the genome of OregS90 shows evidence for expansions of Helitron, LTR *Gypsy* and LTR *Copia* families. On average, 10% of all TEs overlap with genes. Overall, singleton and accessory genes tend to be closer to TEs and contain more often TE insertions than core genes (Additional file 1: Figure S12-S13). The isolates Aus01 and OregS90 have 12.8% and 12.4% of all TEs overlapping with genes, respectively. In addition, Aus01 and OregS90 isolates have 7.4% and 5.4% of all genes that overlap with TEs, respectively (Additional file 1: Figure S14). The composition of TEs inserted into genes reflects the overall TE composition in the genome, with more abundant TEs being more often inserted into genes (Additional file 1: Figure S15). TEs can carry their own regulatory sequences and are often epigenetically silenced by the host. We found that orthogroups comprising a gene within 100 bp distance of a TE show stronger expression variation (∼62% of orthogroups with a coefficient of variation >50%) compared to other orthogroups (∼54% of orthogroups with a coefficient of variation >50%) (Additional file 1: Figure S16-S17). We also found that different TE superfamilies have contrasting effects on gene expression, with longer TEs having more drastic effects (Figure 5D). On average, genes with an inserted TE have lower expression levels (log10 CPM ∼1.7-fold) and a higher coefficient of variation (log10 CPM ∼2-fold) compared to genes without an inserted TE (Additional file 1: Figure S18).

### TE transcription correlates with relative frequency across isolates

Class I TEs replicate through an RNA intermediate and class II through a DNA intermediate. Nevertheless, class II TEs can also transcribe into RNA. To gain insights into the mechanisms of proliferation, we analysed the relative abundance of TE-derived transcripts across all genomes. The highly repetitive nature of TEs typically prevents expression quantification at the individual copy level. Hence, we focused on normalized TE expression across all copies. Overall, more than 70% of the TE families have non-zero transcription levels. This is consistent with recent findings of pervasive transcription of TEs in the *Z. tritici* genome under nutrient stress and during infection (68). We find that the largest TE family, an unclassified LTR identified as RLX_LARD_Thrym, was the most transcribed with an average log_10_ CPM ∼ 4.2 (Figure 6A). An unclassified DTX-MITE is the second most transcribed TE with an average log_10_ CPM ∼ 3.6 followed by an unclassified TE (XXX_*Hermione* with an average log_10_ CPM ∼ 3.4). At the superfamily level, LINEs have the highest expression overall followed by the aggregation of unclassified TEs (Figure 6B). Retroelements are more transcribed than DNA transposons (average log_10_ CPM ∼2 and 1.2, respectively).

**Figure 6:**
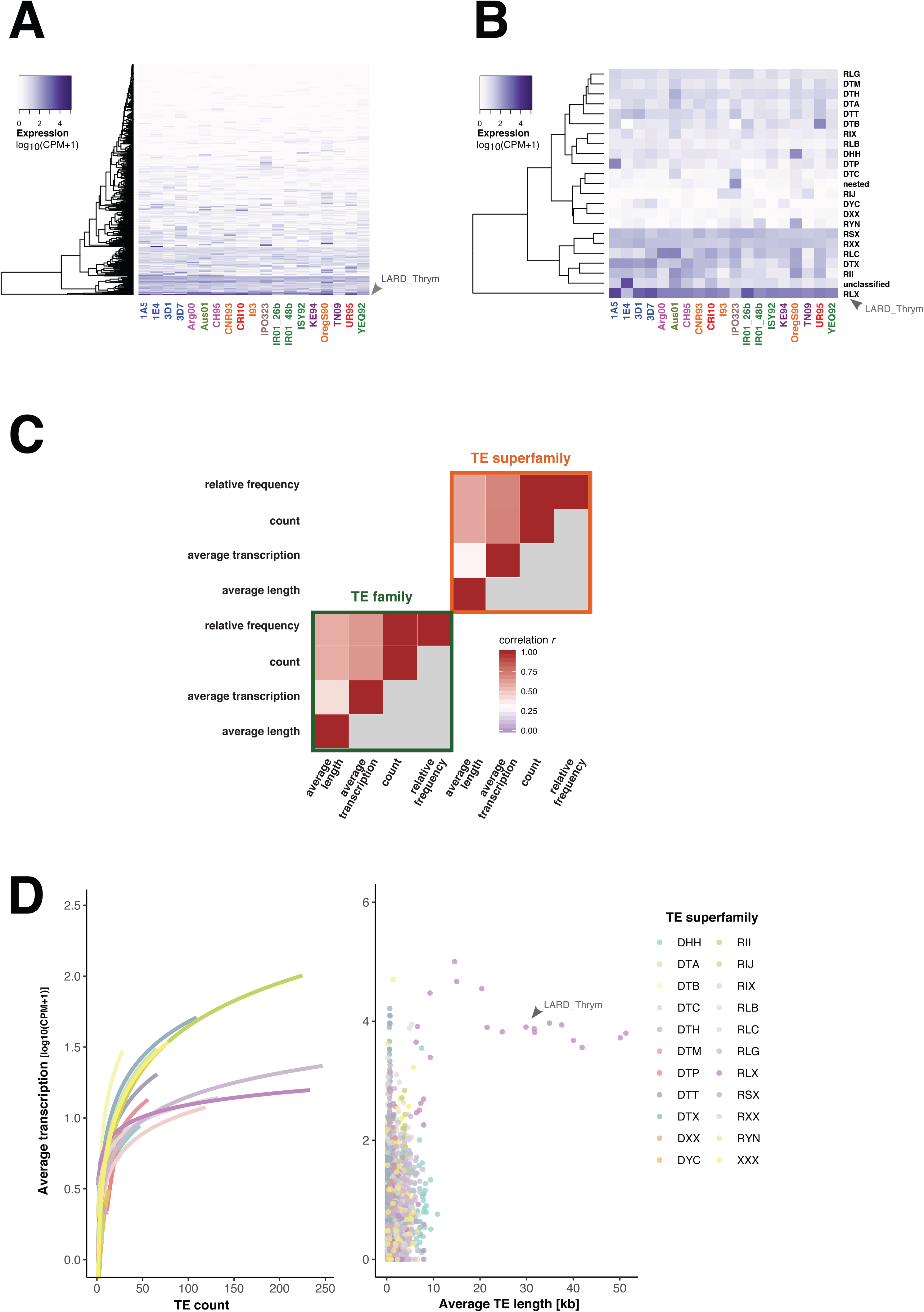
Transcriptional activity of transposable elements (TEs). **A**. TE family transcription levels across all 19 genomes expressed as *log_10_* (CPM +1). **B**. Average transcription levels of TE superfamilies across all genomes expressed as average *log_10_* (CPM +1). **C**. Spearman correlation matrix of four TE metrics including counts, relative frequencies, average length and transcription both at the level of TE families and superfamilies. **D**. Variation of TE transcription (average *log_10_* (CPM +1)) as a function of TE counts (left panel) or average TE length (right panel). Curves in the left panel show the logarithmic linear regression given by the linear model *log_10_* (CPM+1) *∼ log_10_* (TE count). The highly expressed LARD_Thrym family (RLX) is highlighted using arrows (panels A, B, and D).

To understand TE expression dynamics across the pangenome, we investigated associations between TE transcription, length and relative frequency (Figure 6C). We found TE transcription to be correlated with TE frequency in the genomes (Spearman’s *r* = 0.49, *p* < 5e-307; Figure 6C) and we found an even stronger correlation at the TE superfamily level (Spearman’s *r* = 0.59, *p* < 5e-40). However, TE transcription is not correlated with TE length at the superfamily level (Spearman’s *r* =0.06, *p >* 2e-1; Figure 6C). Interestingly, the average TE transcription levels are positively correlated with TE count in the genome (Figure 6D). A notable exception is unclassified SINE retroelements. The correlation of TE transcription levels and TE frequency in the genome strongly suggests that transcriptional activity contributed to recent TE expansions in the genome.

## Discussion

We established a global pangenome of a major fungal wheat pathogen based on the assembly and analysis of 19 high-quality genomes. *Z. tritici* segregates major chromosomal rearrangements affecting both the more conserved core chromosomes as well as the highly polymorphic accessory chromosomes. The gene content is highly variable among genomes with only 60% of all genes being conserved in the species. Accessory genes encode functions for a wide variety of interactions with both biotic and abiotic environments. An exhaustive map of TEs across all genomes pinpoints transposon-associated genome expansions across geographic regions.

We showed that the *Z. tritici* pangenome is expansive with ∼40% accessory orthogroups. Compared to a previous construction of the *Z. tritici* pangenome based on genomes from a much narrower geographic breadth (27), we used more relaxed criteria to assign genes into orthogroups. Based on the tendency to assign more divergent gene variants into the same orthogroup, we recovered a total of 911 orthogroups with at least one paralog compared to only 76 identified previously. The number of paralogs remains low compared to species with larger genomes that retained more paralogs of gene duplication events (28). A likely constraint on gene duplication is the genomic defence mechanism that introduces repeat-induced point (RIP) mutations (45). Although these defences evolved to suppress transpositional activity of TEs, they can also affect genome evolution by targeting gene duplicates (45, 46). Recent sequencing efforts oriented around important crop species reported impressively large accessory genome proportions (47–49). However, nearly all eukaryotic pangenomes are partially based on short-read assemblies that challenge the resolution of segregating gene variants within a species. With the conservative estimate of ∼24% non-reference orthogroups, the *Z. tritici* accessory genome is the largest reported for a fungal species to date (∼40% of the pangenome). This falls outside the upper range of comparative analyses of human fungal pathogens and *S. cerevisiae,* where estimates of the accessory genome ranged from 10-20% (26). However, bacterial accessory genomes can range from 0 to 95% of the total pangenome (3). The effective population size of a species, its lifestyle, and niche heterogeneity are main factors influencing bacterial pangenome sizes (4). Similar to bacteria, the effective population size is likely to be the major factor maintaining a large accessory genome in *Z. tritici*. Previous studies identified *Z. tritici* as a highly polymorphic species with a rapid decay in linkage disequilibrium, high SNP densities and high recombination rates (32, 35). As a consequence, the pathogen likely retains significant functional variation within populations as long as the variation is nearly neutral.

Bacterial and fungal genomes show clear functional compartmentalization between core and accessory genes (4, 26). In fungi, core orthogroups are enriched for housekeeping functions in contrast to an enrichment for antimicrobial resistance and pathogenicity factors among accessory genes (27). Here we show that genes encoding carbohydrate-active enzymes (CAZymes) are highly conserved within the species. CAZymes are involved in the degradation of the host cell wall and other storage compounds (50, 51). Strong conservation of the content in CAZymes may reflect a fundamental adaptation to wheat as a host plant. This contrasts with generalist pathogens, which often evolved larger CAZyme repertoires (52). In contrast to CAZymes, secondary metabolite gene clusters show substantial presence-absence variation within the species. Fungi produce highly diverse secondary metabolites that play a role during various life cycle stages, but often have poorly understood functions (53). Plant pathogens were also shown to depend on secondary metabolite production for full virulence (54). Hence, variation in secondary metabolite production may underlie variation in virulence. Species from the genus *Aspergillus* produce a large diversity of secondary metabolites for which the gene clusters often segregate presence-absence (55, 56). The *Z. tritici* pangenome was constructed from isolates coming from six different continents and a wide array of agricultural environments. Hence, differences in secondary metabolite production capacity may reflect local adaptation and trade-offs that balance the cost of metabolite production. Virulence of *Z. tritici* is thought to be largely governed by gene-for-gene interactions (57). In such interactions effector proteins either promote disease or are recognized by the host and trigger resistance (19). A gene encoding a recognized effector should therefore be rapidly eliminated from the species gene pool. *Z. tritici* populations responded rapidly to selection on effector gene loci by either mutating, deleting or silencing genes (21,33,34). Our global pangenome analysis significantly expands our understanding of effector gene diversification. We identified 652 orthogroups encoding predicted effector functions of which 63% are accessory orthogroups. Accessory effector genes may be involved in arms races with strong selection driving the gain or loss of individual effector genes in populations. As a contrast, we identified 45 conserved and highly expressed effectors genes potentially encoding indispensable pathogenicity functions.

Ultimate mechanisms promoting intra-specific diversity in genome structure may include large population sizes and niche complexity, however the proximate mechanisms generating such diversification are poorly understood. TEs can be key drivers generating structural variation (58, 59) and *Z. tritici* readily undergoes TE-mediated chromosomal rearrangements during meiosis (38, 39). Here we show that *Z. tritici* genomes contain 16-24% TEs, with the overall proportion of TEs accounting for ∼70% of the intraspecific genome size variation. Hence, TEs are key drivers of genome evolution in this species. Among the most drastic chromosomal rearrangements, we detected a significantly shorter chromosome 7 homolog. The longer homolog was hypothesized to have originated from a fusion with an accessory chromosome based on evidence from large scale epigenetic remodelling (60). Our analysis likely identified the ancestral variant prior to the suspected chromosomal fusion event. Hence, the species retained two major chromosomal variants of a core chromosome.

TEs are often implicated in gene copy number variation through duplication or pseudogenisation events suggesting that TEs directly contribute to pangenome diversification. We show that specific *Gypsy* and *Helitron* elements were integrated into genes generating highly paralogous orthogroups. These orthogroups may underlie recent expansions of specific TEs in the genomes of Australian and Oregon isolates. The *Helitron* element is among the most transcribed TEs in the Oregon isolate, suggesting a high potential for new transpositions. In contrast, the *Gypsy* element is only weakly transcribed in the Australian isolate, suggesting that this TE has become deactivated by genomic defences. In addition to transpositional activity causing loss-of-function mutations in genes, TEs can also contribute to genome expansions (61). We found a strong correlation of TE content and genome size across the pangenome suggesting that TEs are the primary drivers of genome expansions. Because he pathogen was only recently introduced to regions outside of Europe and Asia, genome size variation among geographic regions may have originated from population bottlenecks such as founder events. As an example, populations in Australia underwent a significant founder event during the recent colonization of the continent from Europe (62). Hence, our observation of an expanded Australian genome may be causally linked to this bottleneck. Genome expansions may also be triggered by TE mobilisation. Stressors such as host defences during infection cause substantial TE de-repression across the *Z. tritici* genome (68). Taken together, TE dynamics and large effective population sizes likely constitute the proximate and ultimate drivers of pangenome size evolution. Understanding the birth and death cycles of gene functions in such evolving pangenomes will help address major questions related to crop-pathogen co-evolution.

## Methods

### High molecular-weight DNA extraction and single molecule real-time (SMRT) sequencing

Origin and year of sampling of all the isolates are described in Supplementary TableS3. High-molecular-weight DNA was extracted from lyophilized spores following a modified version of a cetyltrimethylammonium bromide (CTAB) protocol developed for plant tissue described in (40). Briefly, ∼100 mg of lyophilized spores were crushed with a mortar and transferred to a phenol-chloroform-isoamyl alcohol solution. The supernatant was centrifuged and the pellet resuspended twice in fresh phenol-chloroform-isoamyl alcohol. The resulting pellet was then washed three times and resuspended in 100 µl of sterile water. For each isolate, PacBio SMRTbell libraries were prepared using between 15 μg and 30 μg of high molecular-weight DNA. Sequencing was performed on a PacBio Sequel instrument at the Functional Genomics Center, Zürich, Switzerland.

### Complete genome assemblies

We largely followed the pipeline described in (64). In summary, raw PacBio sequencing reads were assembled using *Canu* v1.7.1 (65). All assemblies were performed with an estimated genome size of 39.678 Mb (--genomeSize). Two corrected error rates (--correctedErrorRate 0.045 and 0.039) and minimal read length (--minReadLength 500 and 5000) parameters were tested and the most contiguous chromosome-level assemblies were retained for further analysis based on reference alignment. The scaffolding was quality-controlled by inspecting genome-wide dot plots against previously assembled and validated genomes for reference. For each isolate, raw reads were aligned to the newly assembled genome using *pbalign* v0.3.1 from Pacific Biosciences suite (https://github.com/PacificBiosciences/pbalign) to inspect potential mis-assemblies. The assemblies were polished twice using PacBio reads mapped back to the new assembly using the software Arrow v2.2.2 from the Pacific Biosciences suite with default settings (https://github.com/PacificBiosciences/GenomicConsensus) and chromosome-level assemblies were performed using Ragout v2.1.1 and the IPO323 isolate as a reference (66).

### RNA extraction, library preparation, sequencing and quantification

For isolates 1A5, 1E4, 3D1 and 3D7, RNA sequencing experiments on minimal media were performed by (67, 68). Raw reads were retrieved from the NCBI Short Read Archive accession number SRP077418. Similarly, the 15 additional fungal isolates (Additional file 2: Table S3) were grown in YSB media (10g sucrose + 10g yeast extract per liter) and then 10e5 cells were inoculated on liquid minimal media without a carbon source (69) for 7-10 days prior to extraction to reach identical growth stages as for the previous RNA sequencing experiments. RNA was extracted using a NucleoSpin® RNA Plant kit following the manufacturer’s instructions. Library preparation was carried out according to the Illumina TruSeq Stranded mRNA Library Prep protocol with unique indexes for each sample. Single-end 100-bp sequencing was performed on a HiSeq 4000 at the iGE3 platform in Geneva, Switzerland. RNA-seq reads were first filtered using Trimmomatic v0.38 (70) using the following parameters: ILLUMINACLIP:TruSeq3-SE.fa: 2:30:10 LEADING:10 TRAILING:10 SLIDINGWINDOW:5:10 MINLEN: 50, and then aligned to the corresponding genome assembly using STAR v2.6.0a (71) allowing for multiple read mapping (parameters set as --outFilterMultimapNmax 100 --winAnchorMultimapNmax 200 --outFilterMismatchNmax 3). We used HTSeq-count v0.11.2 (72) with -s reverse and –m union parameters to recover counts per feature (joint counting of reads in genes and TEs). We calculated normalized feature counts expressed as counts per million, which accounts for library size, using the EdgeR package v3.24.3 (73). We restricted our analyses to features with a count per million >1.

### Gene prediction and genome annotation

We used the gene prediction pipeline BRAKER v2.1 to predict genes in the 14 newly assembled genomes (74–81). BRAKER combines coding sequence and intron hints based on the mapping of conserved protein sequences and introns identified in RNA-seq data, respectively. The above described RNA-seq datasets were joined with predicted protein sequences from the reference isolate IPO323 (28) and used to predict gene features and guide splice site mapping. RNA alignment files were generated with HISAT2 v2.1.0 using the --rna-strandness R option (82). The resulting bam files were provided to BRAKER (--bam option) together with mapped IPO323 reference proteins (--prot_seq option) to generate gene predictions for each assembled genome using the --alternatives-from-evidence=false --prg=gth --etpmode --fungus parameters. Orthologous genes were identified using protein sequences from all 19 isolates and Orthofinder v2.1.2 with default parameters (83, 84).

### TE consensus identification, classification and annotation

To obtain consensus sequences for TE families, individual runs of RepeatModeler were performed on the 19 complete genomes in addition to the genome of *Z. pseudotritici* (85). The classification was based on the GIRI Repbase using RepeatMasker (86, 87). In order to finalize the classification of TE consensus sequences, we used WICKERsoft (88). The 19 complete genomes were screened for copies of consensus sequences with blastn filtering for sequence identity of > 80% on >80% of the length of the sequence (89). Flanks of 300 bp were added and new multiple sequence alignments were performed using ClustalW (90). Boundaries were visually inspected and trimmed if necessary. Consensus sequences were classified according to the presence and type of terminal repeats and homology of encoded proteins using hints from blastx on NCBI. Consensus sequences were renamed according to a three-letter classification system (91).

A second round of annotation was performed based on predicted protein sequences of TE superfamilies from other fungal species. Here again, the 19 complete genomes were screened for a protein sequence of each superfamily using tblastn. Blast hits were filtered for a minimal alignment size of 80 bp and sequence similarity >35%. Flanks of 3’000 bp or more both up and downstream of the sequence were then added. Hits were pairwise compared with dotplots using dotter and grouped into families based on visual inspection (92). Finally, multiple sequence alignments were performed with ClustalW to construct consensus sequences and the consensus sequences were renamed according to the three-letter system (91).

A third round of annotation of the 19 complete genomes was done to identify four groups of short non-autonomous TEs. LTR-Finder was used to screen for LARDs (LArge Retrotransposon Derivates) and TRIMs (Terminal Repeat retrotransposons In Miniature) with the filters -d 2001 -D 6000 -l 30 -L 5000 and -d 30 -D 2000 -l 30 -L 500 respectively. MITE-Tracker was used to screen for MITEs (Miniature Inverted-repeat Transposable Elements) and SINE-Finder in Sine-Scan to screen for SINEs (Short Interspersed Nuclear Elements) (93–98). For each detected LARD, TRIM and SINE, consensus sequences were created as described above and duplicates excluded. All genome assemblies were then annotated with the curated consensus sequences using RepeatMasker with a cut-off value of 250 and ignored simple repeats as well as low complexity regions. Annotated elements shorter than 100 bp were filtered out, and adjacent identical TEs overlapping by more than 100 bp were merged. Different TE families overlapping by more than 100 bp were considered as *nested* insertions and were renamed accordingly. Identical elements separated by less than 200 bp indicative of putative interrupted elements were grouped into a single element using minimal start and maximal stop positions. TEs overlapping ≥ 1 bp with genes were recovered using the *bedtools* v2.27.1 suite and the *overlap* function (99). Correlations were calculated in RStudio version 1.1.453 using Spearman’s coefficient for pairwise complete observations and statistics were inferred with the *psych* package using the Holm correction method (100).

### Functional annotation of predicted genes

Protein functions were predicted for all gene models using InterProScan v 5.31-70.0 (101) adding - goterms -iprlookup and -pathway information. Secretion peptides and transmembrane domains (TM) were identified using SignalP v 4.1 and Phobius (102, 103). The secretome was defined as the set of proteins with a signal peptide but no TM as predicted by either SignalP and Phobius. Putative effectors were identified among the set of secreted proteins using EffectorP v 2.0 (104). Carbohydrate-active enzymes (CAZymes) were identified using dbCAN2 release 7.0 server (105, 106) with the three tools HMMER, DIAMOND and Hotpep (107–109). Proteins were classified as a CAZyme if predicted by each of the three tools. We searched for secondary metabolite gene clusters using the online version 4 of antiSMASH (110). Genes belonging to an identified cluster were annotated as “biosynthetic”, “biosynthetic-additional”, “transport”, “regulatory” or “other”. Gene clusters mapping at a conserved, orthologous locus shared by two or more isolate were considered as syntenic.

### Declarations

Ethics approval and consent to participate: n/a

Consent for publication: n/a

#### Availability of data and materials

The genome assembly and annotation for new genome assemblies are available at the European Nucleotide Archive (http://www.ebi.ac.uk/ena) under the BioProject PRJEB33986. The RNA-sequencing raw sequencing data was deposited at the NCBI Short Read Archive under the accession number PRJNA559981.

#### Authors’ contributions

TB and DC conceived the study; TB and UO performed analyses; LA and BAM provided datasets and strains; BAM and DC provided funding; TB and DC wrote the manuscript. All authors read and approved the final manuscript.

## Acknowledgements

We are grateful for helpful comments by Simone Fouché on a previous version of this manuscript. Data generated for this manuscript was obtained in collaboration with the Genetic Diversity Centre (GDC), ETH Zurich and the Functional Genomics Center Zurich (FGCZ).

## Additional Files

### Additional File 1: Supplementary_Figures (.docx)

**Figure S1:** Integrative Genomics Viewer screenshot of PacBio reads aligned back to the YEQ92 genome assembly at the fusion locus between chromosomes 15 and 16.

**Figure S2:** Percent identity given by the multiple protein sequence alignment for each orthogroup. Protein sequences were aligned using *mafft* and alignment identity was extracted with the *easel alistat* tool from Eddy Rivas (https://github.com/EddyRivasLab/easel).

**Figure S3:** Presence-absence heatmap of the orthogroups assigned to secondary metabolite gene clusters. Each line stands for an orthogroup. Syntenic gene clusters are numbered from 1 to 39. Orthogroups including more than one gene per cluster are shown in darker blue (0-3 genes were found assigned to an orthogroup in this analysis).

**Figure S4:** Evolutionary origins of the secondary metabolite genes clusters. We performed *blast* searches using all annotated biosynthetic and biosynthetic-additional proteins as query against four closely related sister species of *Zymoseptoria tritici*. The heatmap shows the percent identity of the top hit found in the four sister species for each of the 147 genes encoding biosynthetic functions in putative gene clusters. The isolates Zpa63, Zp13, Zb87 and Za17 correspond to the species *Z. passerinii*, *Z. pseudotritici*, *Z. brevis* and *Z. ardabiliae* respectively.

**Figure S5:** Gene expression across pangenome categories. Gene expression is shown as log10 values of counts per million reads + 1 because non-expressed genes are also shown.

**Figure S6:** Single-gene core orthogroups heatmap following hierarchical clustering based on Euclidian distances. Gene expression is shown as the log10 values of counts per million reads + 1 as non-expressed genes are also shown.

**Figure S7:** GC-content across transposable element family consensus sequences.

**Figure S8:** Transposable element (TE) content correlated with genome length for both core and accessory chromosomes. The proportion of TEs was calculated as the percentage of chromosome length in bp.

**Figure S9:** The proportion of the genome covered by genes correlated with total genome size.

**Figure S10:** Heatmap of average transposable element size (*log10* of the average length in bp).

**Figure S11:** Heatmap of average transposable element size summarized by superfamily (*log10* of the average TE superfamily length in bp).

**Figure S12:** Distance to closest transposable element across pangenome categories given as log_10_ values of the distance in base pairs.

**Figure S13:** Proportion of pangenome categories overlapping with transposable elements (TE). All features with at least 1 bp overlap with a TE sequence were considered.

**Figure S14:** Proportion of overlapping genes in blue and transposable elements (TE) in grey. All features with at least 1 bp overlap with a TE sequence were considered.

**Figure S15:** Genome-wide transposable element (TE) superfamily frequencies correlated with the proportion of TEs overlapping genes. Proportions are given for each TE superfamily (color code) and each of the 19 isolates.

**Figure S16:** Frequency of orthogroups showing high (> 50%) and low (< 50%) expression coefficient of variation. Only orthogroups were distinguished whether at least one gene of the orthogroup was located within 100 bp of a transposable element or not.

**Figure S17:** Gene expression as a function of its distance to the closest transposable element (TE). The relationship is shown for each of the TE superfamilies across the 19 isolates. Gene expression is given by the log10 values of normalized counts per millions reads +1 as genes showing zero expression are also included.

**Figure S18:** Gene expression of genes overlapping at least one base pair with a transposable element (“yes”) compared to genes not overlapping (“no”). Gene expression is given by *log10* values of counts per million reads.

### Additional File 2: Supplementary_Tables (.xlsx)

**Table S1:** List of all identified orthogroups and pangenome categorization.

**Table S2:** List of the genes encoding Major Facilitator Superfamily domains (IPR036259)

**Table S3:** Summary table of the analyzed isolates.

## References

1. Tettelin H, Riley D, Cattuto C, Medini D. Comparative genomics: the bacterial pan-genome. Curr Opin Microbiol [Internet]. 2008 Oct [cited 2019 Jul 19];11(5):472–7. Available from: http://www.ncbi.nlm.nih.gov/pubmed/19086349

2. Ramasamy D, Mishra AK, Lagier J-C, Padhmanabhan R, Rossi M, Sentausa E, et al. A polyphasic strategy incorporating genomic data for the taxonomic description of novel bacterial species. Int J Syst Evol Microbiol [Internet]. 2014 Feb 1 [cited 2019 Jul 17];64(Pt 2):384–91. Available from: http://www.ncbi.nlm.nih.gov/pubmed/24505076

3. Rouli L, Merhej V, Fournier P-E, Raoult D. The bacterial pangenome as a new tool for analysing pathogenic bacteria. New microbes new Infect [Internet]. 2015 Sep [cited 2019 Jul 17];7:72–85. Available from: http://www.ncbi.nlm.nih.gov/pubmed/26442149

4. McInerney JO, McNally A, O’Connell MJ. Why prokaryotes have pangenomes. Nat Microbiol [Internet]. 2017 Apr 28 [cited 2019 Aug 13];2(4):17040. Available from: http://www.nature.com/articles/nmicrobiol201740

5. Lefébure T, Pavinski Bitar PD, Suzuki H, Stanhope MJ. Evolutionary Dynamics of Complete Campylobacter Pan-Genomes and the Bacterial Species Concept. Genome Biol Evol [Internet]. 2010 Jan 1 [cited 2019 Jul 17];2:646–55. Available from: http://www.ncbi.nlm.nih.gov/pubmed/20688752

6. Sánchez-Vallet A, Fouché S, Fudal I, Hartmann FE, Soyer JL, Tellier A, et al. The Genome Biology of Effector Gene Evolution in Filamentous Plant Pathogens. Annu Rev Phytopathol [Internet]. 2018 Aug 25 [cited 2019 Aug 13];56(1):21–40. Available from: https://www.annualreviews.org/doi/10.1146/annurev-phyto-080516-035303

7. Jackson RW, Vinatzer B, Arnold DL, Dorus S, Murillo J. The influence of the accessory genome on bacterial pathogen evolution. Mob Genet Elements [Internet]. 2011 May [cited 2019 Aug 26];1(1):55–65. Available from: http://www.ncbi.nlm.nih.gov/pubmed/22016845

8. Wu Y, Zaiden N, Cao B. The Core- and Pan-Genomic Analyses of the Genus Comamonas: From Environmental Adaptation to Potential Virulence. Front Microbiol [Internet]. 2018 Dec 12 [cited 2019 Aug 26];9:3096. Available from: https://www.frontiersin.org/article/10.3389/fmicb.2018.03096/full

9. Sánchez-Vallet A, Hartmann FE, Marcel TC, Croll D. Nature’s genetic screens: using genome-wide association studies for effector discovery. Mol Plant Pathol [Internet]. 2018 Jan [cited 2019 Aug 26];19(1):3–6. Available from: http://www.ncbi.nlm.nih.gov/pubmed/29226559

10. Marschall T, Marz M, Abeel T, Dijkstra L, Dutilh BE, Ghaffaari A, et al. Computational pangenomics: status, promises and challenges. Brief Bioinform [Internet]. 2016 Oct 21 [cited 2019 Aug 26];19(1):bbw089. Available from: https://academic.oup.com/bib/article-lookup/doi/10.1093/bib/bbw089

11. Schrider DR, Hahn MW. Gene copy-number polymorphism in nature. Proc R Soc B Biol Sci [Internet]. 2010 Nov 7 [cited 2019 Aug 13];277(1698):3213–21. Available from: http://www.ncbi.nlm.nih.gov/pubmed/20591863

12. Brynildsrud O, Gulla S, Feil EJ, Nørstebø SF, Rhodes LD. Identifying copy number variation of the dominant virulence factors msa and p22 within genomes of the fish pathogen Renibacterium salmoninarum. Microb genomics [Internet]. 2016 [cited 2019 Jul 19];2(4):e000055. Available from: http://www.ncbi.nlm.nih.gov/pubmed/28348850

13. Plissonneau C, Daverdin G, Ollivier B, Blaise F, Degrave A, Fudal I, et al. A game of hide and seek between avirulence genes *AvrLm4-7* and *AvrLm3* in *Leptosphaeria maculans*. New Phytol [Internet]. 2016 Mar [cited 2017 Sep 13];209(4):1613–24. Available from: http://www.ncbi.nlm.nih.gov/pubmed/26592855

14. Hartmann FE, Rodríguez de la Vega RC, Brandenburg J-T, Carpentier F, Giraud T. Gene Presence–Absence Polymorphism in Castrating Anther-Smut Fungi: Recent Gene Gains and Phylogeographic Structure. Van De Peer Y, editor. Genome Biol Evol [Internet]. 2018 May 1 [cited 2019 Jul 19];10(5):1298–314. Available from: https://academic.oup.com/gbe/article/10/5/1298/4990910

15. Araki H, Tian D, Goss EM, Jakob K, Halldorsdottir SS, Kreitman M, et al. Presence/absence polymorphism for alternative pathogenicity islands in Pseudomonas viridiflava, a pathogen of Arabidopsis. Pnas. 2006;103(15):5887–92.

16. Wit De PJGM, Mehrabi R, Burg Van den HA, Stergiopoulos I. Fungal effector proteins: past, present and future. Mol Plant Pathol [Internet]. 2009 Nov [cited 2017 Sep 18];10(6):735–47. Available from: http://www.ncbi.nlm.nih.gov/pubmed/19849781

17. Lo Presti L, Lanver D, Schweizer G, Tanaka S, Liang L, Tollot M, et al. Fungal Effectors and Plant Susceptibility. Annu Rev Plant Biol [Internet]. 2015 Apr 29 [cited 2017 Sep 13];66(1):513–45. Available from: http://www.ncbi.nlm.nih.gov/pubmed/25923844

18. Toruño TY, Stergiopoulos I, Coaker G. Plant-Pathogen Effectors: Cellular Probes Interfering with Plant Defenses in Spatial and Temporal Manners. Annu Rev Phytopathol [Internet]. 2016 Aug 4 [cited 2019 Aug 26];54(1):419–41. Available from: http://www.annualreviews.org/doi/10.1146/annurev-phyto-080615-100204

19. Jones JDG, Dangl JL. The plant immune system. Nature [Internet]. 2006 Nov 16 [cited 2014 Jul 10];444(7117):323–9. Available from: http://www.ncbi.nlm.nih.gov/pubmed/17108957

20. Yoshida K, Saunders DGO, Mitsuoka C, Natsume S, Kosugi S, Saitoh H, et al. Host specialization of the blast fungus Magnaporthe oryzae is associated with dynamic gain and loss of genes linked to transposable elements. BMC Genomics [Internet]. 2016 Dec 18 [cited 2019 Jul 19];17(1):370. Available from: http://bmcgenomics.biomedcentral.com/articles/10.1186/s12864-016-2690-6

21. Hartmann FE, Croll D. Distinct Trajectories of Massive Recent Gene Gains and Losses in Populations of a Microbial Eukaryotic Pathogen. Mol Biol Evol [Internet]. 2017 Jul 21 [cited 2017 Sep 13];127(19):1–18. Available from: https://academic.oup.com/mbe/article-lookup/doi/10.1093/molbev/msx208

22. Faino L, Seidl MF, Shi-Kunne X, Pauper M, Van Den Berg GCM, Wittenberg AHJ, et al. Transposons passively and actively contribute to evolution of the two-speed genome of a fungal pathogen. Genome Res. 2016;26(8):1091–100.

23. Sperschneider J, Gardiner DM, Thatcher LF, Lyons R, Singh KB, Manners JM, et al. Genome-Wide Analysis in Three Fusarium Pathogens Identifies Rapidly Evolving Chromosomes and Genes Associated with Pathogenicity. Genome Biol Evol [Internet]. 2015 May 19 [cited 2017 Sep 13];7(6):1613–27. Available from: http://www.ncbi.nlm.nih.gov/pubmed/25994930

24. Dong S, Raffaele S, Kamoun S. The two-speed genomes of filamentous pathogens: waltz with plants. Curr Opin Genet Dev [Internet]. 2015 Dec [cited 2019 Jul 19];35:57–65. Available from: http://www.ncbi.nlm.nih.gov/pubmed/26451981

25. Fouché S, Plissonneau C, Croll D. The birth and death of effectors in rapidly evolving filamentous pathogen genomes. Curr Opin Microbiol [Internet]. 2018 Dec [cited 2019 Aug 13];46:34–42. Available from: http://www.ncbi.nlm.nih.gov/pubmed/29455143

26. McCarthy CGP, Fitzpatrick DA. Pan-genome analyses of model fungal species. Microb genomics [Internet]. 2019 [cited 2019 Jul 19];5(2). Available from: http://www.ncbi.nlm.nih.gov/pubmed/30714895

27. Plissonneau C, Hartmann FE, Croll D. Pangenome analyses of the wheat pathogen *Zymoseptoria tritici* reveal the structural basis of a highly plastic eukaryotic genome. BMC Biol [Internet]. 2018 Jan 11 [cited 2018 Feb 6];16(1):5. Available from: https://bmcbiol.biomedcentral.com/articles/10.1186/s12915-017-0457-4

28. Goodwin SB, Ben M’Barek S, Dhillon B, Wittenberg AHJ, Crane CF, Hane JK, et al. Finished Genome of the Fungal Wheat Pathogen Mycosphaerella graminicola Reveals Dispensome Structure, Chromosome Plasticity, and Stealth Pathogenesis. Malik HS, editor. PLoS Genet [Internet]. 2011 Jun 9 [cited 2019 Jun 6];7(6):e1002070. Available from: https://dx.plos.org/10.1371/journal.pgen.1002070

29. Cools HJ, Fraaije BA. Are azole fungicides losing ground against Septoria wheat disease? Resistance mechanisms inMycosphaerella graminicola. Pest Manag Sci [Internet]. 2008 Jul 1 [cited 2019 Aug 13];64(7):681–4. Available from: http://doi.wiley.com/10.1002/ps.1568

30. Blake JJ, Gosling P, Fraaije BA, Burnett FJ, Knight SM, Kildea S, et al. Changes in field dose-response curves for demethylation inhibitor (DMI) and quinone outside inhibitor (QoI) fungicides against *Zymoseptoria tritici*, related to laboratory sensitivity phenotyping and genotyping assays. Pest Manag Sci [Internet]. 2018 Feb 1 [cited 2019 Aug 13];74(2):302–13. Available from: http://doi.wiley.com/10.1002/ps.4725

31. Lucas JA, Hawkins NJ, Fraaije BA. The Evolution of Fungicide Resistance. In: Advances in applied microbiology [Internet]. 2015 [cited 2019 Aug 13]. p. 29–92. Available from: http://www.ncbi.nlm.nih.gov/pubmed/25596029

32. Hartmann FE, Sánchez-Vallet A, McDonald BA, Croll D. A fungal wheat pathogen evolved host specialization by extensive chromosomal rearrangements. ISME J [Internet]. 2017 May 24 [cited 2017 Sep 13];11(5):1189–204. Available from: http://www.ncbi.nlm.nih.gov/pubmed/28117833

33. Meile L, Croll D, Brunner PC, Plissonneau C, Hartmann FE, McDonald BA, et al. A fungal avirulence factor encoded in a highly plastic genomic region triggers partial resistance to septoria tritici blotch. New Phytol [Internet]. 2018 [cited 2019 Aug 26];219(3):1048–61. Available from: http://www.ncbi.nlm.nih.gov/pubmed/29693722

34. Krishnan P, Meile L, Plissonneau C, Ma X, Hartmann FE, Croll D, et al. Transposable element insertions shape gene regulation and melanin production in a fungal pathogen of wheat. BMC Biol [Internet]. 2018 Dec 16 [cited 2019 Aug 26];16(1):78. Available from: http://www.ncbi.nlm.nih.gov/pubmed/30012138

35. Croll D, Lendenmann MH, Stewart E, McDonald BA. The impact of recombination hotspots on genome evolution of a fungal plant pathogen. Genetics [Internet]. 2015 Nov [cited 2017 Aug 19];201(3):1213–28. Available from: http://www.ncbi.nlm.nih.gov/pubmed/26392286

36. Grandaubert J, Dutheil JY, Stukenbrock EH. The genomic determinants of adaptive evolution in a fungal pathogen. Evol Lett [Internet]. 2019 Jun 1 [cited 2019 Aug 26];3(3):299–312. Available from: http://doi.wiley.com/10.1002/evl3.117

37. Stukenbrock EH, Dutheil JY. Fine-Scale Recombination Maps of Fungal Plant Pathogens Reveal Dynamic Recombination Landscapes and Intragenic Hotspots. Genetics [Internet]. 2018 Mar 1 [cited 2019 Aug 26];208(3):1209–29. Available from: http://www.ncbi.nlm.nih.gov/pubmed/29263029

38. Croll D, Zala M, McDonald BA, Smoot M, Shumway M. Breakage-fusion-bridge Cycles and Large Insertions Contribute to the Rapid Evolution of Accessory Chromosomes in a Fungal Pathogen. Heitman J, editor. PLoS Genet [Internet]. 2013 Jun 13 [cited 2017 Sep 13];9(6):e1003567. Available from: http://dx.plos.org/10.1371/journal.pgen.1003567

39. Fouché S, Plissonneau C, McDonald BA, Croll D. Meiosis Leads to Pervasive Copy-Number Variation and Distorted Inheritance of Accessory Chromosomes of the Wheat Pathogen Zymoseptoria tritici. Genome Biol Evol [Internet]. 2018 [cited 2019 Aug 26];10(6):1416–29. Available from: http://www.ncbi.nlm.nih.gov/pubmed/29850789

40. Plissonneau C, Stürchler A, Croll D. The Evolution of Orphan Regions in Genomes of a Fungal Pathogen of Wheat. MBio [Internet]. 2016 Nov 18 [cited 2019 Feb 22];7(5):e01231–16. Available from: http://www.ncbi.nlm.nih.gov/pubmed/27795389

41. Steinberg G. Cell biology of Zymoseptoria tritici: Pathogen cell organization and wheat infection. Fungal Genet Biol [Internet]. 2015 Jun [cited 2019 Aug 13];79:17–23. Available from: http://www.ncbi.nlm.nih.gov/pubmed/26092785

42. Palma-Guerrero J, Ma X, Torriani SFF, Zala M, Francisco CS, Hartmann FE, et al. Comparative Transcriptome Analyses in *Zymoseptoria tritici* Reveal Significant Differences in Gene Expression Among Strains During Plant Infection. Mol Plant-Microbe Interact [Internet]. 2017 Mar [cited 2017 Sep 13];30(3):231–44. Available from: http://www.ncbi.nlm.nih.gov/pubmed/28121239

43. Steinhauer D, Salat M, Frey R, Mosbach A, Luksch T, Balmer D, et al. A dispensable paralog of succinate dehydrogenase subunit C mediates standing resistance towards a subclass of SDHI fungicides in Zymoseptoria tritici. bioRxiv [Internet]. 2019 Apr 23 [cited 2019 Jul 8];616904. Available from: https://www.biorxiv.org/content/10.1101/616904v1

44. Omrane S, Audéon C, Ignace A, Duplaix C, Aouini L, Kema G, et al. Plasticity of the MFS1 Promoter Leads to Multidrug Resistance in the Wheat Pathogen Zymoseptoria tritici. mSphere [Internet]. 2017 Oct 25 [cited 2019 Sep 12];2(5):e00393-17. Available from: http://www.ncbi.nlm.nih.gov/pubmed/29085913

45. Selker EU. Repeat-Induced Gene Silencing in Fungi. Adv Genet [Internet]. 2002 Jan 1 [cited 2019 Aug 26];46:439–50. Available from: https://www.sciencedirect.com/science/article/abs/pii/S0065266002460166?via%3Dihub

46. Galagan JE, Selker EU. RIP: the evolutionary cost of genome defense. Trends Genet [Internet]. 2004 Sep 1 [cited 2019 Aug 26];20(9):417–23. Available from: http://www.ncbi.nlm.nih.gov/pubmed/15313550

47. Hirsch CN, Foerster JM, Johnson JM, Sekhon RS, Muttoni G, Vaillancourt B, et al. Insights into the maize pan-genome and pan-transcriptome. Plant Cell [Internet]. 2014 Jan 1 [cited 2019 Aug 16];26(1):121–35. Available from: http://www.ncbi.nlm.nih.gov/pubmed/24488960

48. Zhou P, Silverstein KAT, Ramaraj T, Guhlin J, Denny R, Liu J, et al. Exploring structural variation and gene family architecture with De Novo assemblies of 15 Medicago genomes. BMC Genomics [Internet]. 2017 [cited 2019 Aug 16];18(1):261. Available from: http://www.ncbi.nlm.nih.gov/pubmed/28347275

49. Zhao Q, Feng Q, Lu H, Li Y, Wang A, Tian Q, et al. Pan-genome analysis highlights the extent of genomic variation in cultivated and wild rice. Nat Genet [Internet]. 2018 Feb 15 [cited 2019 Jul 19];50(2):278–84. Available from: http://www.nature.com/articles/s41588-018-0041-z

50. Lyu X, Shen C, Fu Y, Xie J, Jiang D, Li G, et al. Comparative genomic and transcriptional analyses of the carbohydrate-active enzymes and secretomes of phytopathogenic fungi reveal their significant roles during infection and development. Sci Rep [Internet]. 2015 Dec 4 [cited 2019 Aug 16];5(1):15565. Available from: http://www.nature.com/articles/srep15565

51. Zerillo MM, Adhikari BN, Hamilton JP, Buell CR, Lévesque CA, Tisserat N. Carbohydrate-Active Enzymes in Pythium and Their Role in Plant Cell Wall and Storage Polysaccharide Degradation. Lespinet O, editor. PLoS One [Internet]. 2013 Sep 12 [cited 2019 Aug 16];8(9):e72572. Available from: https://dx.plos.org/10.1371/journal.pone.0072572

52. Zhao Z, Liu H, Wang C, Xu J-R. Comparative analysis of fungal genomes reveals different plant cell wall degrading capacity in fungi. BMC Genomics [Internet]. 2013 Apr 23 [cited 2017 Sep 18];14(1):274. Available from: http://www.ncbi.nlm.nih.gov/pubmed/23617724

53. Calvo AM, Wilson RA, Bok JW, Keller NP. Relationship between secondary metabolism and fungal development. Microbiol Mol Biol Rev [Internet]. 2002 Sep 1 [cited 2019 Aug 16];66(3):447–59, table of contents. Available from: http://www.ncbi.nlm.nih.gov/pubmed/12208999

54. Pusztahelyi T, Holb IJ, Pócsi I. Secondary metabolites in fungus-plant interactions. Front Plant Sci [Internet]. 2015 [cited 2019 Aug 16];6:573. Available from: http://www.ncbi.nlm.nih.gov/pubmed/26300892

55. Raffa N, Keller NP. A call to arms: Mustering secondary metabolites for success and survival of an opportunistic pathogen. Sheppard DC, editor. PLOS Pathog [Internet]. 2019 Apr 4 [cited 2019 Aug 26];15(4):e1007606. Available from: http://dx.plos.org/10.1371/journal.ppat.1007606

56. Kjærbølling I, Vesth TC, Frisvad JC, Nybo JL, Theobald S, Kuo A, et al. Linking secondary metabolites to gene clusters through genome sequencing of six diverse Aspergillus species. Proc Natl Acad Sci U S A [Internet]. 2018 Jan 23 [cited 2019 Aug 26];115(4):E753–61. Available from: http://www.ncbi.nlm.nih.gov/pubmed/29317534

57. Brown JKM, Chartrain L, Lasserre-Zuber P, Saintenac C. Genetics of resistance to Zymoseptoria tritici and applications to wheat breeding. Fungal Genet Biol [Internet]. 2015 Jun [cited 2019 Aug 26];79:33–41. Available from: http://www.ncbi.nlm.nih.gov/pubmed/26092788

58. Beck CR, Garcia-Perez JL, Badge RM, Moran J V. LINE-1 Elements in Structural Variation and Disease. Annu Rev Genomics Hum Genet [Internet]. 2011 Sep 22 [cited 2019 Feb 20];12(1):187–215. Available from: http://www.annualreviews.org/doi/10.1146/annurev-genom-082509-141802

59. Kim S, Mun S, Kim T, Lee K-H, Kang K, Cho J-Y, et al. Transposable element-mediated structural variation analysis in dog breeds using whole-genome sequencing. Mamm Genome [Internet]. 2019 Aug 15 [cited 2019 Aug 26]; Available from: http://www.ncbi.nlm.nih.gov/pubmed/31414176

60. Schotanus K, Soyer JL, Connolly LR, Grandaubert J, Happel P, Smith KM, et al. Histone modifications rather than the novel regional centromeres of Zymoseptoria tritici distinguish core and accessory chromosomes. Epigenetics Chromatin [Internet]. 2015 Dec 1 [cited 2019 Jul 8];8(1):41. Available from: http://www.epigeneticsandchromatin.com/content/8/1/41

61. Naville M, Henriet S, Warren I, Sumic S, Reeve M, Volff J-N, et al. Massive Changes of Genome Size Driven by Expansions of Non-autonomous Transposable Elements. Curr Biol [Internet]. 2019 Apr 1 [cited 2019 Aug 19];29(7):1161–1168.e6. Available from: https://www.sciencedirect.com/science/article/abs/pii/S0960982219301393

62. Zhan J, Linde CC, Jurgens T, Merz U, Steinebrunner F, McDonald BA. Variation for neutral markers is correlated with variation for quantitative traits in the plant pathogenic fungus Mycosphaerella graminicola. Mol Ecol [Internet]. 2005 Aug 1 [cited 2019 Sep 12];14(9):2683–93. Available from: http://doi.wiley.com/10.1111/j.1365-294X.2005.02638.x

64. Yue J-X, Liti G. Long-read sequencing data analysis for yeasts. Nat Protoc [Internet]. 2018 Jun 3 [cited 2019 Jul 16];13(6):1213–31. Available from: http://www.nature.com/articles/nprot.2018.025

65. Koren S, Walenz BP, Berlin K, Miller JR, Bergman NH, Phillippy AM. Canu: scalable and accurate long-read assembly via adaptive k-mer weighting and repeat separation. Genome Res [Internet]. 2017 [cited 2019 Jul 16];27(5):722–36. Available from: http://www.ncbi.nlm.nih.gov/pubmed/28298431

66. Kolmogorov M, Raney B, Paten B, Pham S. Ragout-a reference-assisted assembly tool for bacterial genomes. Bioinformatics [Internet]. 2014 Jun 15 [cited 2019 Jul 16];30(12):i302–9. Available from: http://www.ncbi.nlm.nih.gov/pubmed/24931998

67. Francisco CS, Ma X, Zwyssig MM, McDonald BA, Palma-Guerrero J. Morphological changes in response to environmental stresses in the fungal plant pathogen Zymoseptoria tritici. Sci Rep [Internet]. 2019 Dec 3 [cited 2019 Oct 2];9(1):9642. Available from: http://www.nature.com/articles/s41598-019-45994-3

68. Fouché S, Badet T, Oggenfuss U, Plissonneau C, Francisco CS, Croll D. Stress-driven transposable element de-repression dynamics and virulence evolution in a fungal pathogen. Arkhipova I, editor. Mol Biol Evol [Internet]. 2019 Sep 25 [cited 2019 Oct 2]; Available from: https://academic.oup.com/mbe/advance-article/doi/10.1093/molbev/msz216/5573762

69. Metzenberg RL. Vogel’s Medium N salts: avoiding the need for ammonium nitrate. Fungal Genet Rep [Internet]. 2003 Dec 1 [cited 2019 Jul 16];50(1):14–14. Available from: https://newprairiepress.org/fgr/vol50/iss1/6

70. Bolger AM, Lohse M, Usadel B. Trimmomatic: a flexible trimmer for Illumina sequence data. Bioinformatics [Internet]. 2014 Aug 1 [cited 2019 Jul 16];30(15):2114–20. Available from: http://www.ncbi.nlm.nih.gov/pubmed/24695404

71. Dobin A, Davis CA, Schlesinger F, Drenkow J, Zaleski C, Jha S, et al. STAR: ultrafast universal RNA-seq aligner. Bioinformatics [Internet]. 2013 Jan 1 [cited 2019 Aug 16];29(1):15–21. Available from: http://www.ncbi.nlm.nih.gov/pubmed/23104886

72. Anders S, Pyl PT, Huber W. HTSeq--a Python framework to work with high-throughput sequencing data. Bioinformatics [Internet]. 2015 Jan 15 [cited 2019 Jul 16];31(2):166–9. Available from: http://www.ncbi.nlm.nih.gov/pubmed/25260700

73. Robinson MD, McCarthy DJ, Smyth GK. edgeR: a Bioconductor package for differential expression analysis of digital gene expression data. Bioinformatics [Internet]. 2010 Jan 1 [cited 2019 Jul 16];26(1):139. Available from: http://www.ncbi.nlm.nih.gov/pubmed/19910308

74. Hoff KJ, Lange S, Lomsadze A, Borodovsky M, Stanke M. BRAKER1: Unsupervised RNA-Seq-Based Genome Annotation with GeneMark-ET and AUGUSTUS: Table 1. Bioinformatics [Internet]. 2016 Mar 1 [cited 2019 Jul 16];32(5):767–9. Available from: http://www.ncbi.nlm.nih.gov/pubmed/26559507

75. Stanke M, Schöffmann O, Morgenstern B, Waack S. Gene prediction in eukaryotes with a generalized hidden Markov model that uses hints from external sources. BMC Bioinformatics [Internet]. 2006 Feb 9 [cited 2019 Jul 16];7:62. Available from: http://www.ncbi.nlm.nih.gov/pubmed/16469098

76. Stanke M, Diekhans M, Baertsch R, Haussler D. Using native and syntenically mapped cDNA alignments to improve de novo gene finding. Bioinformatics [Internet]. 2008 Mar 1 [cited 2019 Jul 16];24(5):637–44. Available from: http://www.ncbi.nlm.nih.gov/pubmed/18218656

77. Altschul SF, Gish W, Miller W, Myers EW, Lipman DJ. Basic local alignment search tool. J Mol Biol [Internet]. 1990 Oct 5 [cited 2014 Jul 10];215(3):403–10. Available from: http://www.ncbi.nlm.nih.gov/pubmed/2231712

78. Camacho C, Coulouris G, Avagyan V, Ma N, Papadopoulos J, Bealer K, et al. BLAST+: architecture and applications. BMC Bioinformatics [Internet]. 2009 Dec 15 [cited 2019 Jul 16];10(1):421. Available from: http://www.biomedcentral.com/1471-2105/10/421

79. Lomsadze A, Burns PD, Borodovsky M. Integration of mapped RNA-Seq reads into automatic training of eukaryotic gene finding algorithm. Nucleic Acids Res [Internet]. 2014 Sep 2 [cited 2019 Jul 16];42(15):e119–e119. Available from: http://www.ncbi.nlm.nih.gov/pubmed/24990371

80. Barnett DW, Garrison EK, Quinlan AR, Stromberg MP, Marth GT. BamTools: a C++ API and toolkit for analyzing and managing BAM files. Bioinformatics [Internet]. 2011 Jun 15 [cited 2019 Jul 16];27(12):1691–2. Available from: https://academic.oup.com/bioinformatics/article-lookup/doi/10.1093/bioinformatics/btr174

81. Li H, Handsaker B, Wysoker A, Fennell T, Ruan J, Homer N, et al. The Sequence Alignment/Map format and SAMtools. Bioinformatics [Internet]. 2009 Aug 15 [cited 2019 Jul 16];25(16):2078–9. Available from: https://academic.oup.com/bioinformatics/article-lookup/doi/10.1093/bioinformatics/btp352

82. Kim D, Langmead B, Salzberg SL. HISAT: a fast spliced aligner with low memory requirements. Nat Methods [Internet]. 2015 Apr 9 [cited 2019 Jul 16];12(4):357–60. Available from: http://www.nature.com/articles/nmeth.3317

83. Emms DM, Kelly S. OrthoFinder: solving fundamental biases in whole genome comparisons dramatically improves orthogroup inference accuracy. Genome Biol [Internet]. 2015 Dec 6 [cited 2019 Jul 16];16(1):157. Available from: http://genomebiology.com/2015/16/1/157

84. Emms DM, Kelly S. OrthoFinder: phylogenetic orthology inference for comparative genomics. bioRxiv [Internet]. 2019 Apr 24 [cited 2019 Jul 16];466201. Available from: https://www.biorxiv.org/content/10.1101/466201v2.full

85. Stukenbrock EH, Jørgensen FG, Zala M, Hansen TT, McDonald BA, Schierup MH. Whole-Genome and Chromosome Evolution Associated with Host Adaptation and Speciation of the Wheat Pathogen Mycosphaerella graminicola. Malik HS, editor. PLoS Genet [Internet]. 2010 Dec 23 [cited 2018 Aug 20];6(12):e1001189. Available from: http://dx.plos.org/10.1371/journal.pgen.1001189

86. Smit, AFA, Hubley, R & Green P. RepeatMasker Open-4.0. [Internet]. 2015. Available from: http://repeatmasker.org

87. Bao W, Kojima KK, Kohany O. Repbase Update, a database of repetitive elements in eukaryotic genomes. Mob DNA [Internet]. 2015 Dec 2 [cited 2019 Jul 16];6(1):11. Available from: http://www.mobilednajournal.com/content/6/1/11

88. Breen J, Wicker T, Kong X, Zhang J, Ma W, Paux E, et al. A highly conserved gene island of three genes on chromosome 3B of hexaploid wheat: diverse gene function and genomic structure maintained in a tightly linked block. BMC Plant Biol [Internet]. 2010 May 27 [cited 2019 Jul 16];10:98. Available from: http://www.ncbi.nlm.nih.gov/pubmed/20507561

89. Altschul SF, Madden TL, Schäffer AA, Zhang J, Zhang Z, Miller W, et al. Gapped BLAST and PSI-BLAST: a new generation of protein database search programs. Nucleic Acids Res. 1997;25(17):3389–402.

90. G Higgins D, M Sharp P. CLUSTAL: a package for performing multiple sequence alignment on a microcomputer. Gene. 1988;73(1):237–44.

91. Wicker T, Sabot F, Hua-Van A, Bennetzen JL, Capy P, Chalhoub B, et al. A unified classification system for eukaryotic transposable elements. Nat Rev Genet. 2007;8(12):973– 82.

92. Sonnhammer ELL, Durbin R. A dot-matrix program with dynamic threshold control suited for genomic DNA and protein sequence analysis. Gene. 1995;167(1–2):Gc1–10.

93. Xu Z, Wang H. LTR-FINDER: An efficient tool for the prediction of full-length LTR retrotransposons. Nucleic Acids Res. 2007;35(SUPPL.2):265–8.

94. Gao D, Li Y, Kim K Do, Abernathy B, Jackson SA. Landscape and evolutionary dynamics of terminal repeat retrotransposons in miniature in plant genomes. Genome Biol [Internet]. 2016 Dec 18 [cited 2019 Jul 16];17(1):7. Available from: http://www.ncbi.nlm.nih.gov/pubmed/26781660

95. Ma B, Li T, Xiang Z, He N. MnTEdb, a collective resource for mulberry transposable elements. Database [Internet]. 2015 Jan 1 [cited 2019 Jul 16];2015. Available from: https://academic.oup.com/database/article/doi/10.1093/database/bav004/2433136

96. Crescente JM, Zavallo D, Helguera M, Vanzetti LS. MITE Tracker: an accurate approach to identify miniature inverted-repeat transposable elements in large genomes. BMC Bioinformatics [Internet]. 2018 Dec 3 [cited 2019 Jul 16];19(1):348. Available from: https://bmcbioinformatics.biomedcentral.com/articles/10.1186/s12859-018-2376-y

97. Mao H, Wang H. SINE_scan: an efficient tool to discover short interspersed nuclear elements (SINEs) in large-scale genomic datasets. Bioinformatics [Internet]. 2017 Jan 6 [cited 2019 Jul 16];33(5):btw718. Available from: http://www.ncbi.nlm.nih.gov/pubmed/28062442

98. Wenke T, Dobel T, Sorensen TR, Junghans H, Weisshaar B, Schmidt T. Targeted Identification of Short Interspersed Nuclear Element Families Shows Their Widespread Existence and Extreme Heterogeneity in Plant Genomes. Plant Cell Online. 2011;23(9):3117–28.

99. Quinlan AR, Hall IM. BEDTools: a flexible suite of utilities for comparing genomic features. Bioinformatics [Internet]. 2010 Mar 15 [cited 2019 Jul 16];26(6):841–2. Available from: https://academic.oup.com/bioinformatics/article-lookup/doi/10.1093/bioinformatics/btq033

100. Revelle WR. psych: Procedures for Personality and Psychological Research [Internet]. 2017 [cited 2019 Aug 16]. Available from: https://www.scholars.northwestern.edu/en/publications/psych-procedures-for-personality-and-psychological-research

101. Jones P, Binns D, Chang H-Y, Fraser M, Li W, McAnulla C, et al. InterProScan 5: genome-scale protein function classification. Bioinformatics [Internet]. 2014 May 1 [cited 2014 Jul 13];30(9):1236–40. Available from: http://www.pubmedcentral.nih.gov/articlerender.fcgi?artid=3998142&tool=pmcentrez&rendertype=abstract

102. Petersen TN, Brunak S, von Heijne G, Nielsen H. SignalP 4.0: discriminating signal peptides from transmembrane regions. Nat Methods [Internet]. 2011 Oct 29 [cited 2019 Jul 16];8(10):785–6. Available from: http://www.ncbi.nlm.nih.gov/pubmed/21959131

103. Käll L, Krogh A, Sonnhammer EL. A Combined Transmembrane Topology and Signal Peptide Prediction Method. J Mol Biol [Internet]. 2004 May 14 [cited 2019 Jul 16];338(5):1027–36. Available from: http://www.ncbi.nlm.nih.gov/pubmed/15111065

104. Sperschneider J, Gardiner DM, Dodds PN, Tini F, Covarelli L, Singh KB, et al. Effector P: predicting fungal effector proteins from secretomes using machine learning. New Phytol [Internet]. 2016 Apr [cited 2019 Jul 16];210(2):743–61. Available from: http://www.ncbi.nlm.nih.gov/pubmed/26680733

105. Zhang H, Yohe T, Huang L, Entwistle S, Wu P, Yang Z, et al. dbCAN2: a meta server for automated carbohydrate-active enzyme annotation. Nucleic Acids Res [Internet]. 2018 Jul 2 [cited 2019 Jul 16];46(W1):W95–101. Available from: http://www.ncbi.nlm.nih.gov/pubmed/29771380

106. Lombard V, Golaconda Ramulu H, Drula E, Coutinho PM, Henrissat B. The carbohydrate-active enzymes database (CAZy) in 2013. Nucleic Acids Res [Internet]. 2014 Jan [cited 2019 Jul 16];42(Database issue):D490-5. Available from: https://academic.oup.com/nar/article-lookup/doi/10.1093/nar/gkt1178

107. Finn RD, Clements J, Eddy SR. HMMER web server: interactive sequence similarity searching. Nucleic Acids Res [Internet]. 2011 Jul [cited 2019 Jul 16];39(Web Server issue):W29–37. Available from: http://www.ncbi.nlm.nih.gov/pubmed/21593126

108. Buchfink B, Xie C, Huson DH. Fast and sensitive protein alignment using DIAMOND. Nat Methods [Internet]. 2015 Jan 17 [cited 2019 Jul 16];12(1):59–60. Available from: http://www.ncbi.nlm.nih.gov/pubmed/25402007

109. Busk PK, Pilgaard B, Lezyk MJ, Meyer AS, Lange L. Homology to peptide pattern for annotation of carbohydrate-active enzymes and prediction of function. BMC Bioinformatics [Internet]. 2017 Dec 12 [cited 2019 Jul 16];18(1):214. Available from: http://www.ncbi.nlm.nih.gov/pubmed/28403817

110. Blin K, Wolf T, Chevrette MG, Lu X, Schwalen CJ, Kautsar SA, et al. antiSMASH 4.0-improvements in chemistry prediction and gene cluster boundary identification. Nucleic Acids Res [Internet]. 2017 [cited 2019 Jul 16];45(W1):W36–41. Available from: http://www.ncbi.nlm.nih.gov/pubmed/28460038

