## Additional file 1 for "A 19-isolate reference-quality global pangenome for the fungal wheat pathogen *Zymoseptoria tritici*"

**Supplementary Figures**

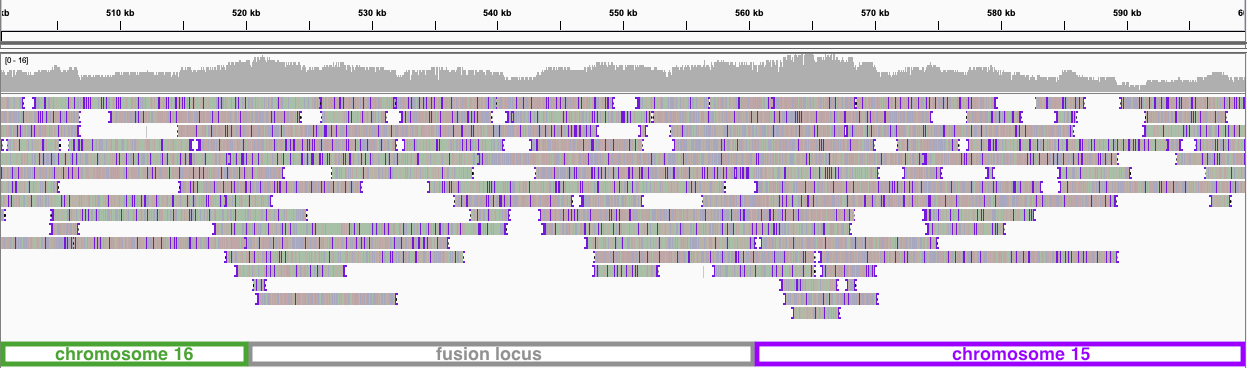

**Figure S1:** Integrative Genomics Viewer screenshot of PacBio reads aligned back to the YEQ92 genome assembly at the fusion locus between chromosomes 15 and 16.

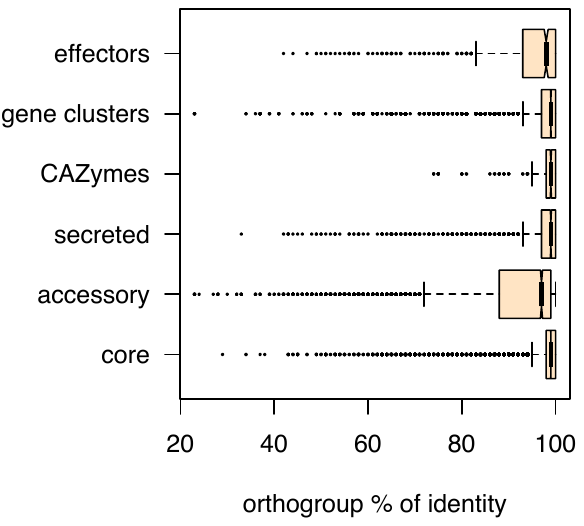

**Figure S2:** Percent identity given by the multiple protein sequence alignment for each orthogroup. Protein sequences were aligned using *mafft* and alignment identity was extracted with the *easel alistat* tool from Eddy Rivas (https://github.com/EddyRivasLab/easel).

**Figure S3:** Presence-absence heatmap of the orthogroups assigned to secondary metabolite gene clusters. Each line stands for an orthogroup. Syntenic gene clusters are numbered from 1 to 39. Orthogroups including more than one gene per cluster are shown in darker blue (0-3 genes were found assigned to an orthogroup in this analysis).

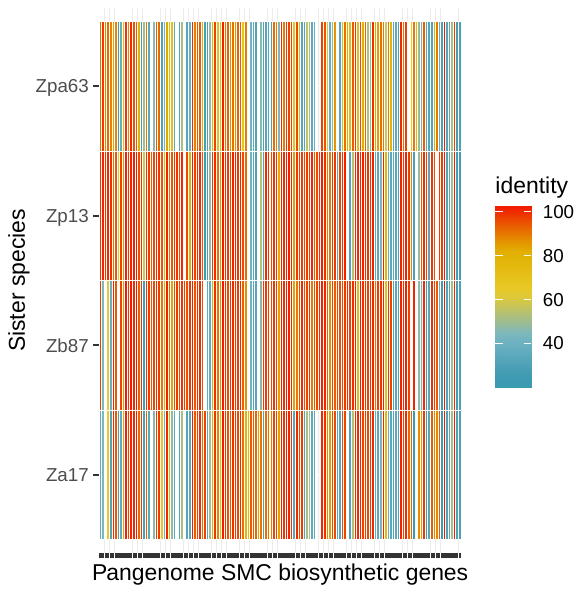

**Figure S4:** Evolutionary origins of the secondary metabolite genes clusters. We performed *blast* searches using all annotated biosynthetic and biosynthetic-additional proteins as query against four closely related sister species of *Zymoseptoria tritici*. The heatmap shows the percent identity of the top hit found in the four sister species for each of the 147 genes encoding biosynthetic functions in putative gene clusters. The isolates Zpa63, Zp13, Zb87 and Za17 correspond to the species *Z. passerinii*, *Z. pseudotritici*, *Z. brevis* and *Z. ardabiliae* respectively.

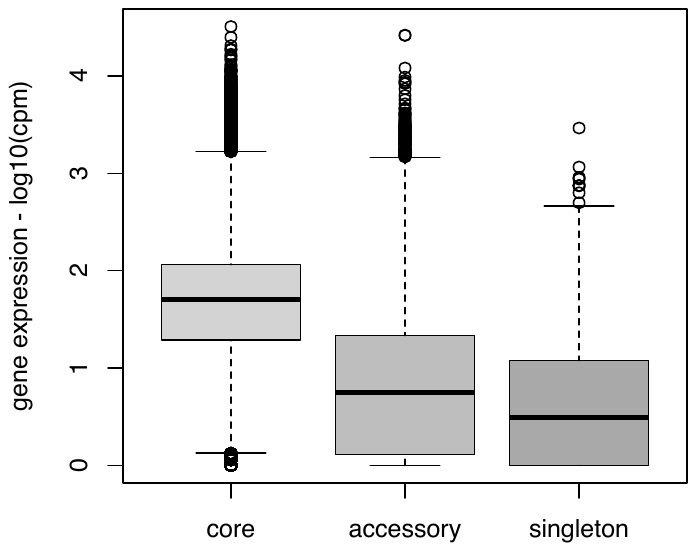

**Figure S5:** Gene expression across pangenome categories. Gene expression is shown as log10 values of counts per million reads + 1 because non-expressed genes are also shown.

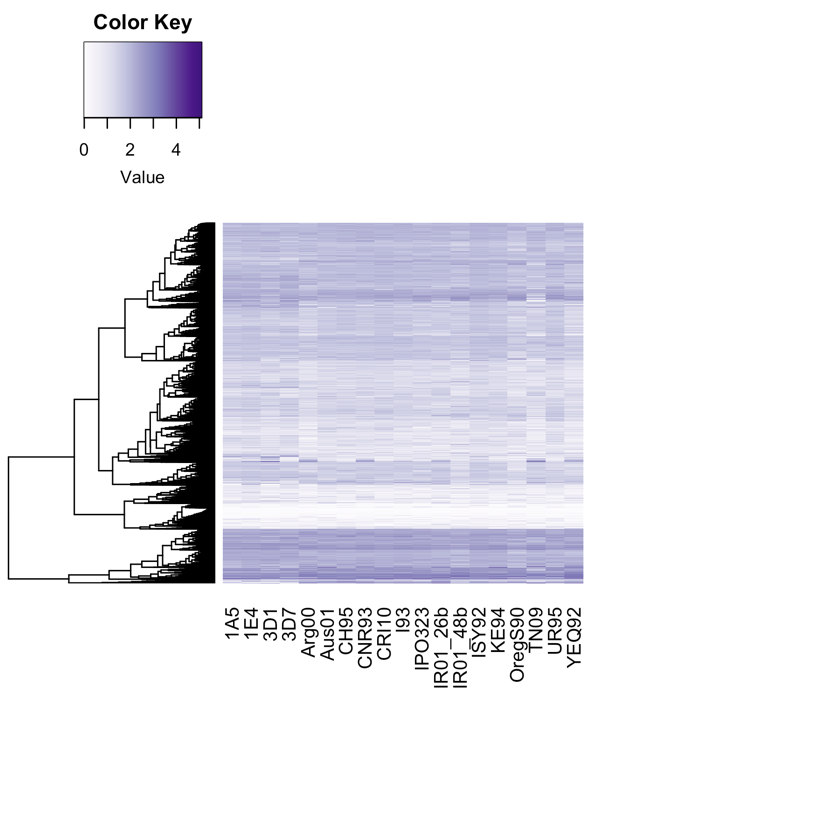

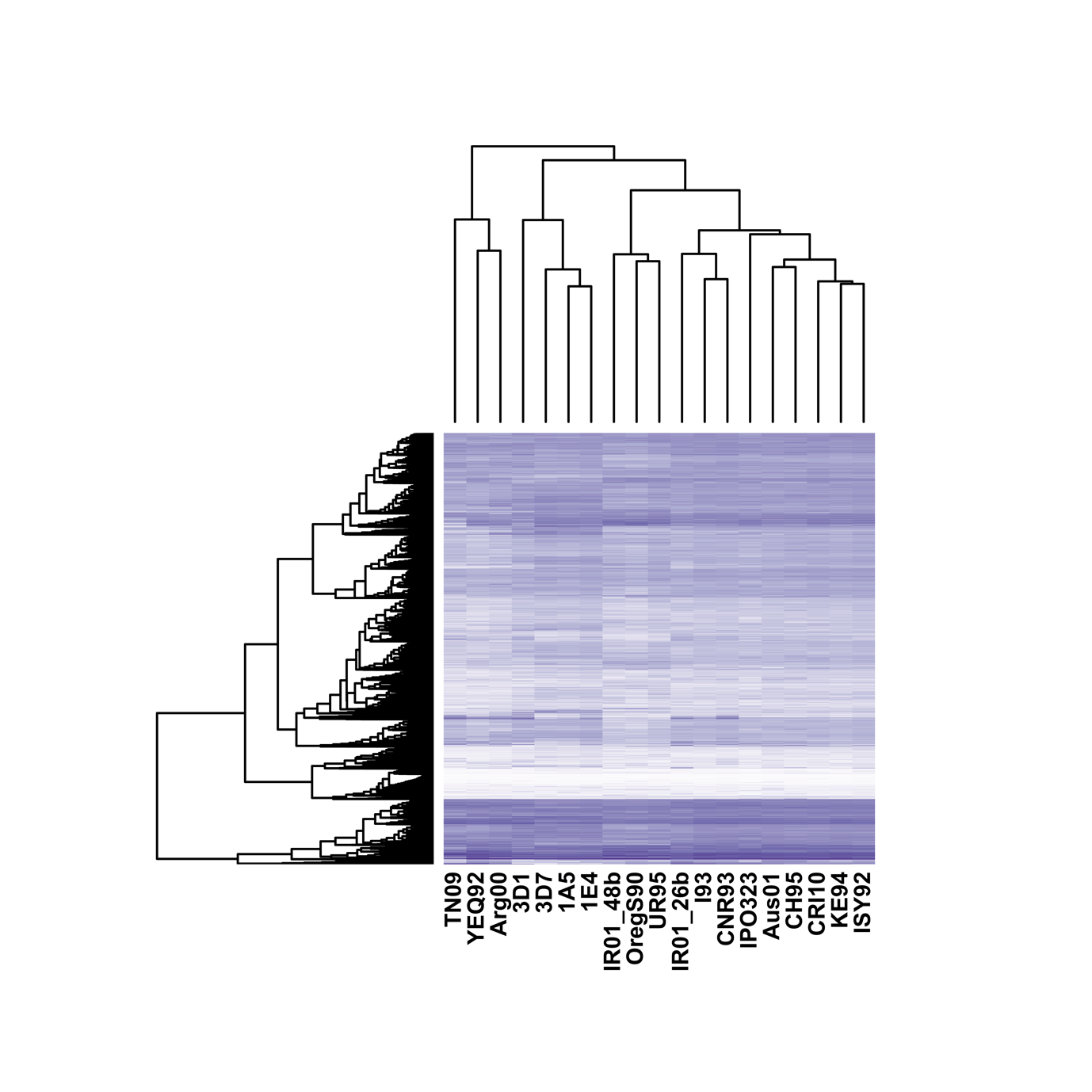

**Figure S6:** Single-gene core orthogroups heatmap following hierarchical clustering based on Euclidian distances. Gene expression is shown as the log10 values of counts per million reads + 1 as non-expressed genes are also shown.

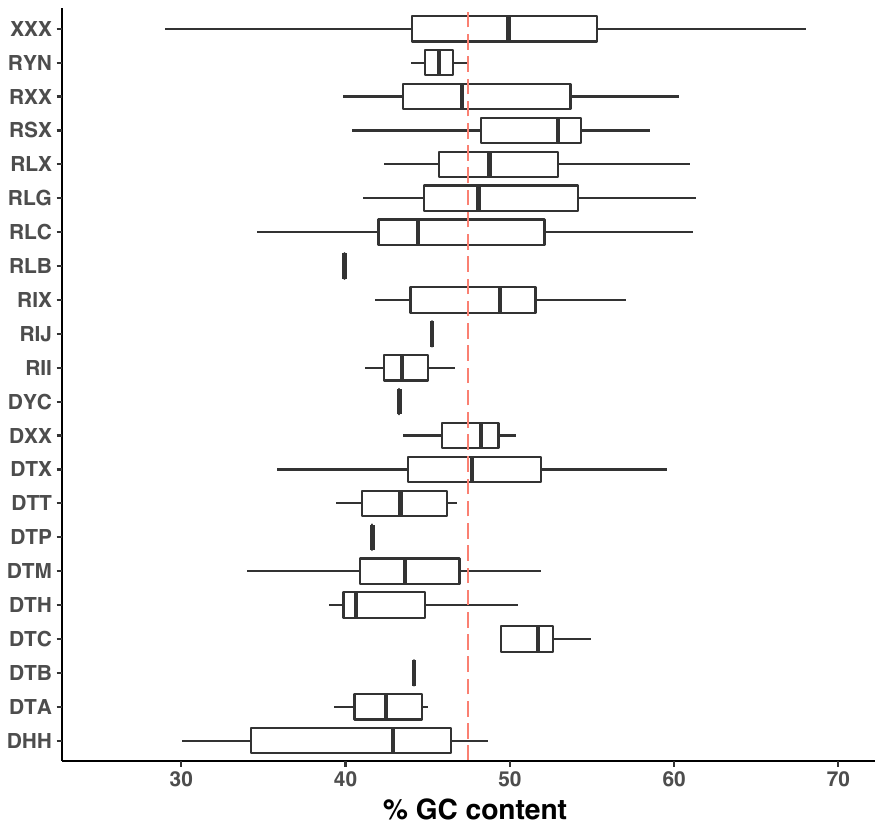

**Figure S7:** GC-content across transposable element family consensus sequences.

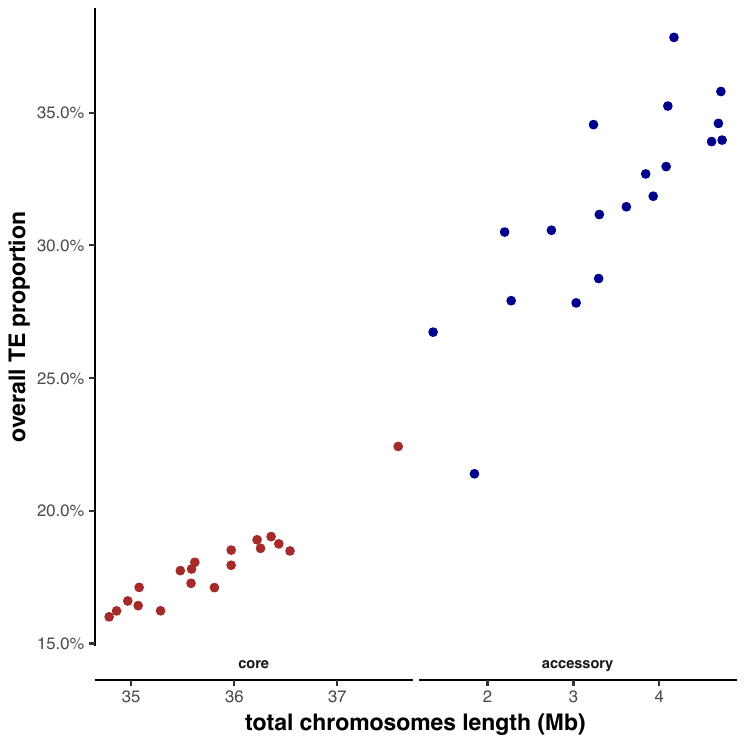

**Figure S8:** Transposable element (TE) content correlated with genome length for both core and accessory chromosomes. The proportion of TEs was calculated as the percentage of chromosome length in bp.

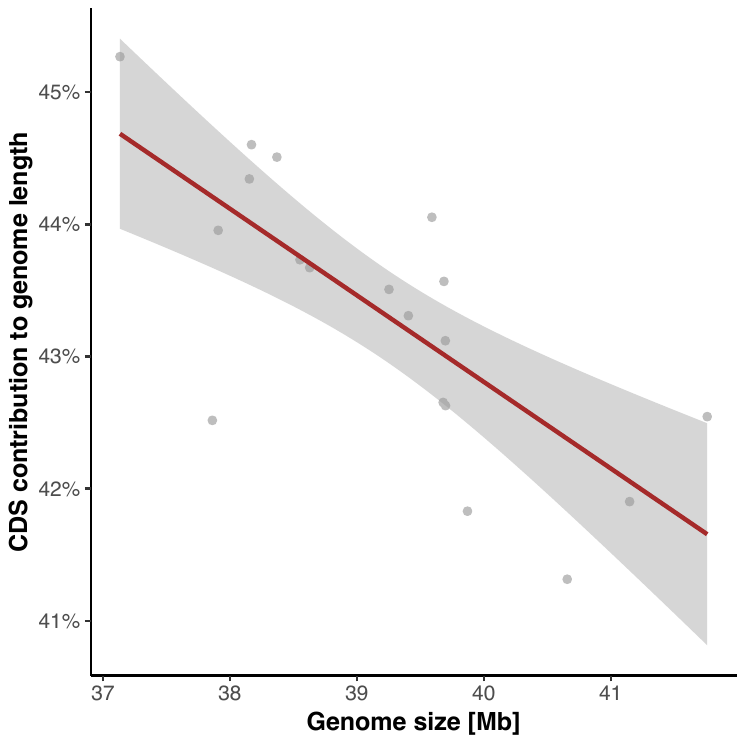

**Figure S9:** The proportion of the genome covered by genes correlated with total genome size.

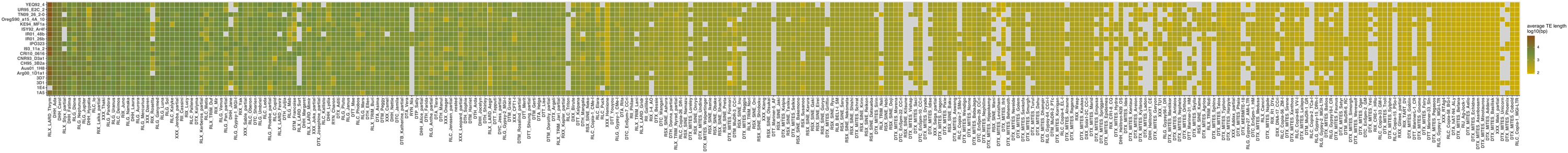

**Figure S10:** Heatmap of average transposable element size (*log10* of the average length in bp).

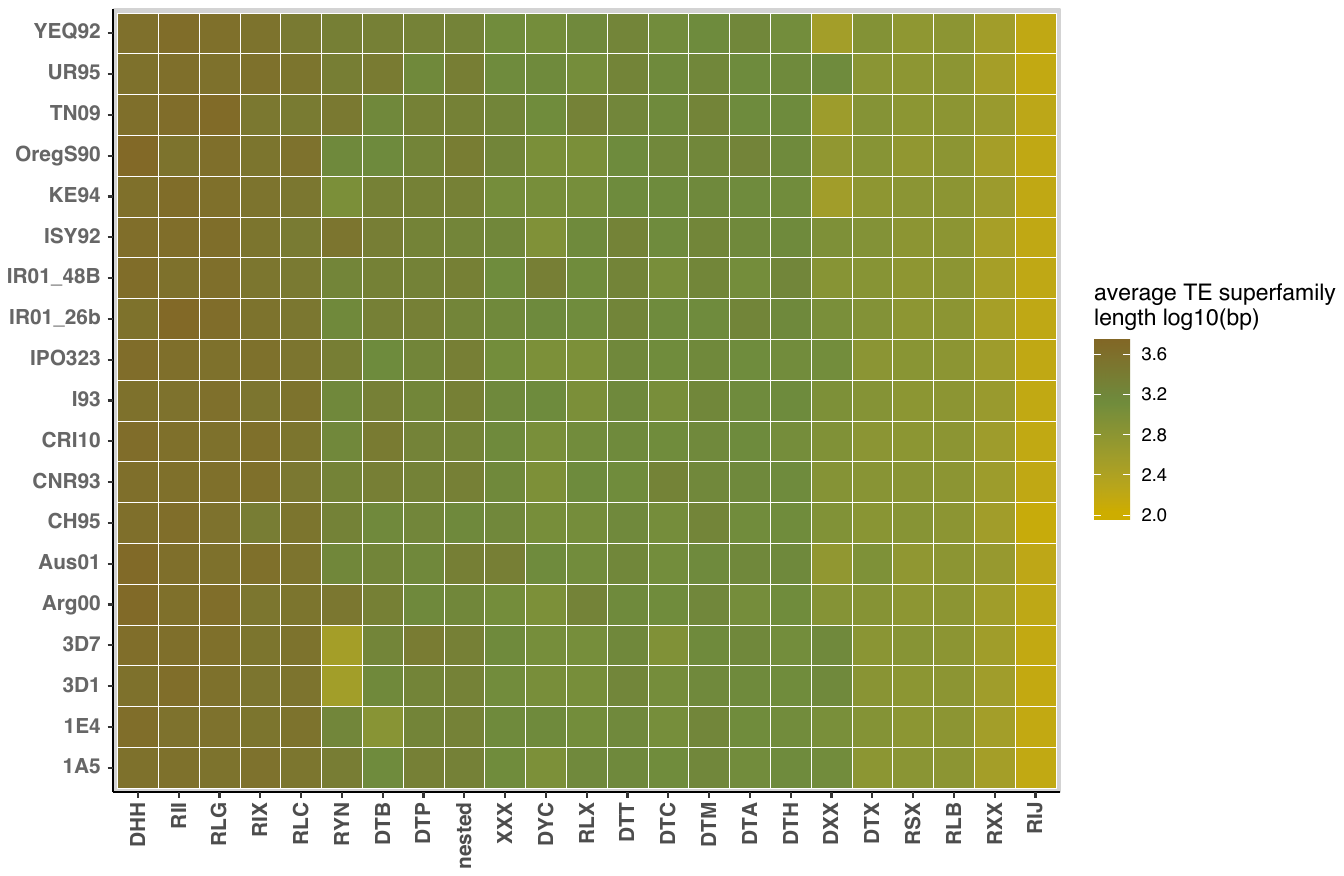

**Figure S11:** Heatmap of average transposable element size summarized by superfamily (*log10* of the average TE superfamily length in bp).

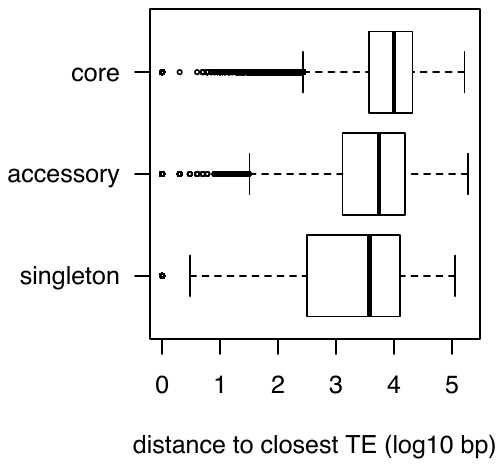

**Figure S12:** Distance to closest transposable element across pangenome categories given as log_10_ values of the distance in base pairs.

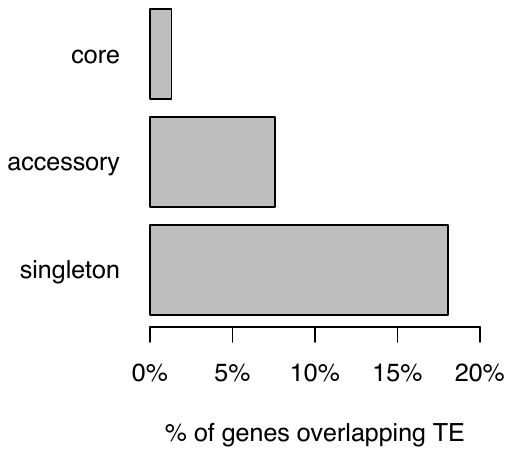

**Figure S13:** Proportion of pangenome categories overlapping with transposable elements (TE). All features with at least 1 bp overlap with a TE sequence were considered.

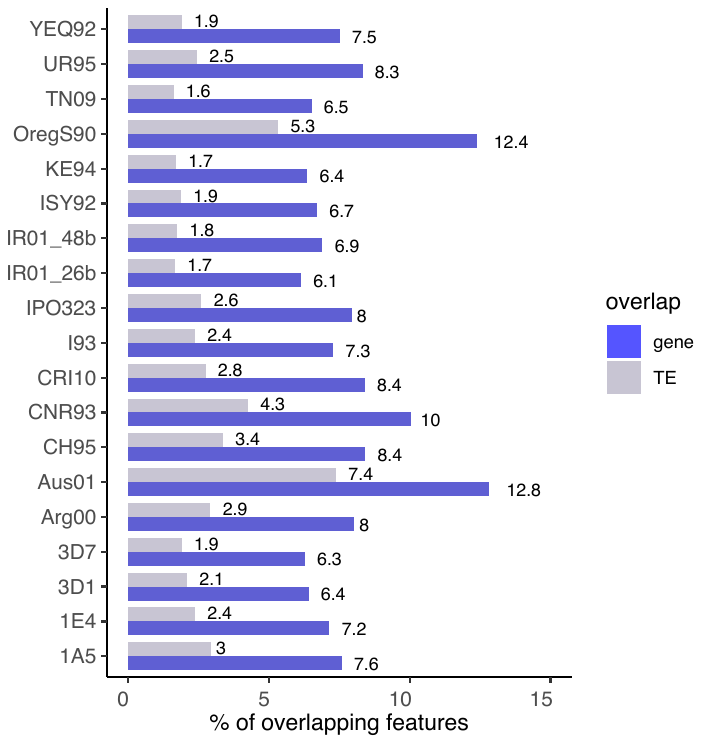

**Figure S14:** Proportion of overlapping genes in blue and transposable elements (TE) in grey. All features with at least 1 bp overlap with a TE sequence were considered.

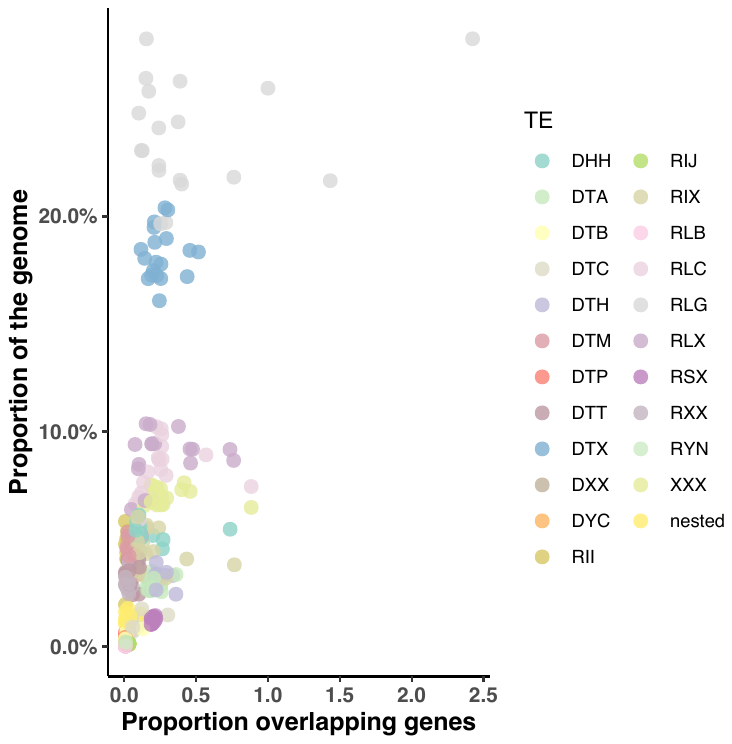

**Figure S15:** Genome-wide transposable element (TE) superfamily frequencies correlated with the proportion of TEs overlapping genes. Proportions are given for each TE superfamily (color code) and each of the 19 isolates.

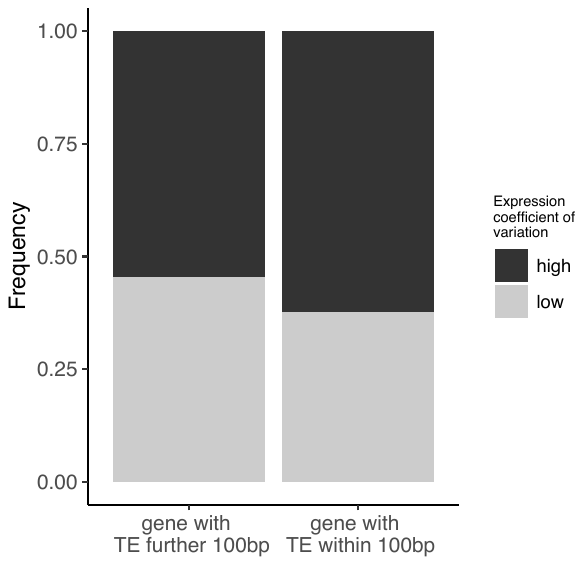

**Figure S16:** Frequency of orthogroups showing high (> 50%) and low (< 50%) expression coefficient of variation. Only orthogroups were distinguished whether at least one gene of the orthogroup was located within 100 bp of a transposable element or not.

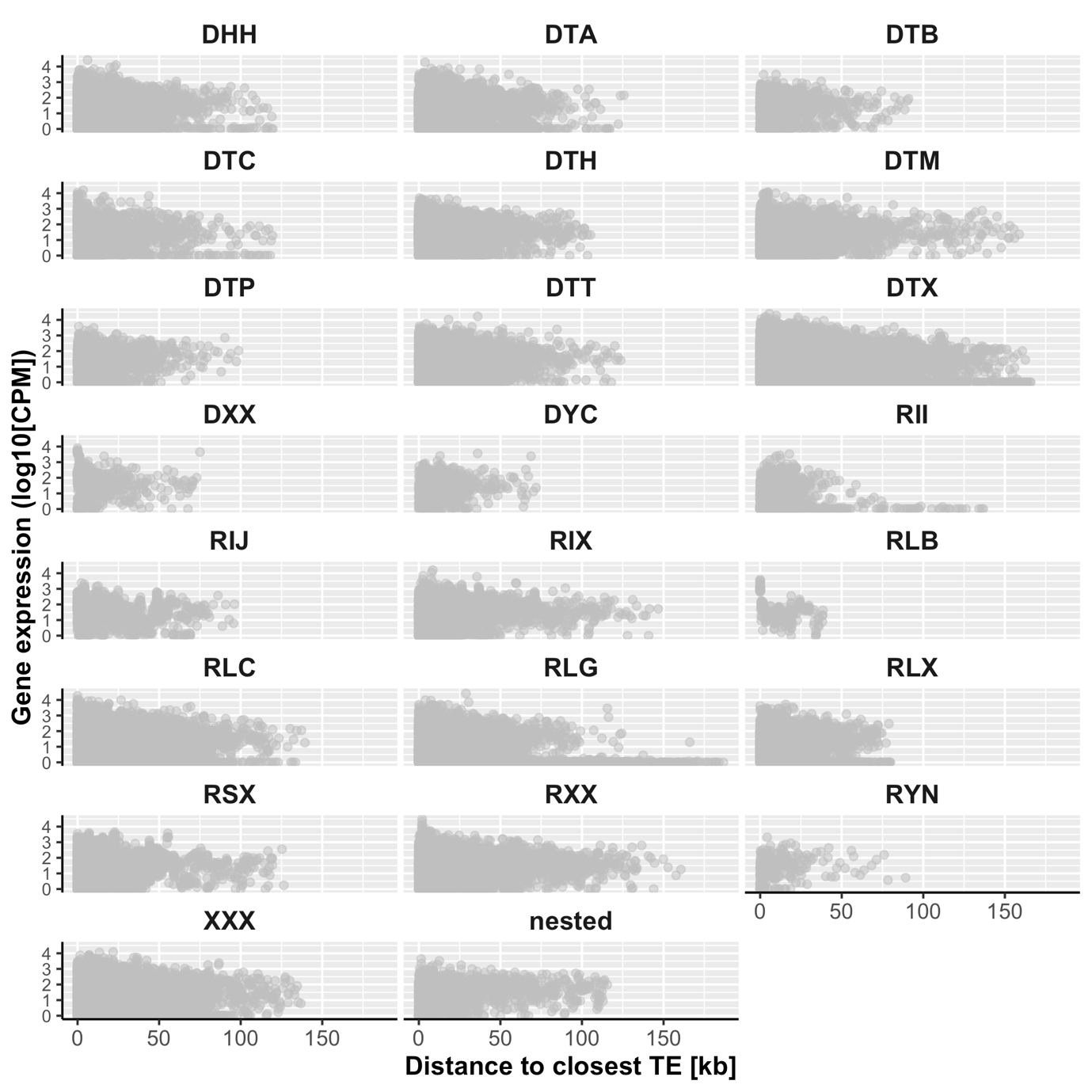

**Figure S17:** Gene expression as a function of its distance to the closest transposable element (TE). The relationship is shown for each of the TE superfamilies across the 19 isolates. Gene expression is given by the log10 values of normalized counts per millions reads +1 as genes showing zero expression are also included.

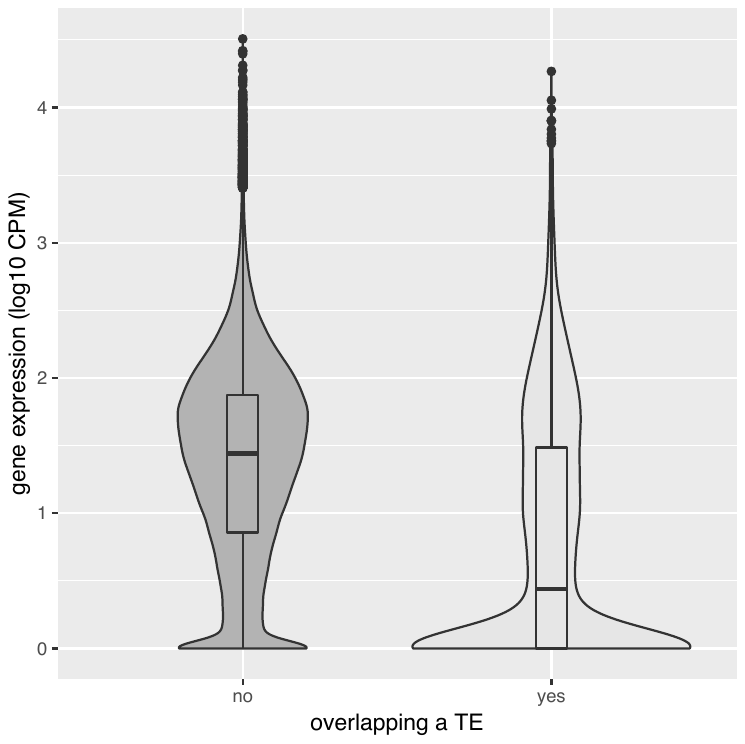

**Figure S18:** Gene expression of genes overlapping at least one base pair with a transposable element (“yes”) compared to genes not overlapping (“no”). Gene expression is given by *log10* values of counts per million reads.
